# Tardigrade CAHS Proteins Act as Molecular Swiss Army Knives to Mediate Desiccation Tolerance Through Multiple Mechanisms

**DOI:** 10.1101/2021.08.16.456555

**Authors:** Cherie S. Hesgrove, Kenny H. Nguyen, Sourav Biswas, Charles A. Childs, KC Shraddha, Bryan X. Medina, Vladimir Alvarado, Feng Yu, Shahar Sukenik, Marco Malferrari, Francesco Francia, Giovanni Venturoli, Erik W. Martin, Alex S. Holehouse, Thomas C. Boothby

**Affiliations:** Molecular Biology Department, University of Wyoming, Laramie, WY, USA; Chemical Engineering Department, University of Wyoming, Laramie, WY USA; Quantitative Systems Biology Program, University of California, Merced, CA, USA; Department of Chemistry and Biochemistry, University of California, Merced, CA, USA; Dipartimento di Chimica “Giacomo Ciamician”, Università di Bologna, via Selmi 2, I-40126 Bologna, Italy; Laboratorio di Biochimica e Biofisica Molecolare, Dipartimento di Farmacia e Biotecnologie, FaBiT, Università di Bologna, via Irnerio 42, I-40126 Bologna, Italy; Consorzio Nazionale Interuniversitario per le Scienze Fisiche della Materia (CNISM), c/o Dipartimento di Fisica e Astronomia (DIFA), Università di Bologna, via Irnerio 46, I-40126 Bologna, Italy; Department of Structural Biology, St. Jude Children’s Research Hospital, Memphis, TN, USA; Department of Biochemistry and Molecular Biophysics, Washington University School of Medicine, St. Louis, MO, USA; Center for Science and Engineering of Living Systems (CSELS), Washington University in St. Louis, St. Louis, MO, USA

## Abstract

Tardigrades, also known as water bears, make up a phylum of small but extremely hardy animals, renowned for their ability to survive extreme stresses, including desiccation. How tardigrades survive desiccation is one of the enduring mysteries of animal physiology. Here we show that CAHS D, an intrinsically disordered protein belonging to a unique family of proteins possessed only by tardigrades, undergoes a liquid-to-gel phase transition in a concentration dependent manner. Unlike other gelling proteins, such as gelatin, our data support a mechanism in which gel formation of CAHS D is driven by intermolecular β-β interactions. We find that gel formation corresponds with strong coordination of water and slowing of water diffusion. The degree of water coordination correlates with the ability of CAHS D to protect lactate dehydrogenase from unfolding when dried. This implies that the mechanism for unfolding protection can be attributed to a combination of hydration and slowed molecular motion. Conversely, rapid diffusion leading to efficient molecular shielding appears to be the predominant mechanism preventing protein aggregation. Our study demonstrates that distinct mechanisms are required for holistic protection during desiccation, and that protectants, such as CAHS D, can act as molecular ‘Swiss Army Knives’ capable of providing protection through several different mechanisms simultaneously.

## Introduction

Anhydrobiosis, the ability to survive near-complete water loss, is an intriguing trait found in all kingdoms of life (Crowe and Clegg, 1973). Extreme drying can impart a number of stresses on biological systems (Boothby, 2019; Boothby and Pielak, 2017; Hesgrove and Boothby, 2020), with protein dysfunctions being a major set of common perturbations (Boothby, 2019; Boothby and Pielak, 2017; Hesgrove and Boothby, 2020). The two prevalent, and non-mutually exclusive, forms of protein dysfunction during desiccation are protein unfolding and protein aggregation (Hesgrove and Boothby, 2020).

Many desiccation-tolerant organisms protect their cells from drying induced damage by accumulating non-reducing sugars, such as sucrose (Yathisha et al., 2020) or trehalose (Erkut et al., 2011; Laskowska and Kuczyńska-Wiśnik, 2020; Mitsumasu et al., 2010; Tapia et al., 2015; Tapia and Koshland, 2014). The enrichment of disaccharides was long thought to be a universal feature of desiccation tolerance. However, several robustly anhydrobiotic organisms, such as tardigrades and rotifers, do not accumulate high levels of sugars during drying (Boothby et al., 2017; Hengherr et al., 2008; Lapinski and Tunnacliffe, 2003). Instead, these animals use a diverse array of intrinsically disordered proteins (IDPs) to provide adaptive protection against desiccation (Boothby et al., 2017; Boothby and Pielak, 2017; Denekamp et al., 2010; Hesgrove and Boothby, 2020; Piszkiewicz et al., 2019; Tripathi, 2012; Tripathi et al., 2012).

One example of stress tolerant IDPs are Cytoplasmic Abundant Heat Soluble (CAHS) proteins, which are employed by tardigrades to survive desiccation (Boothby et al., 2017; Boothby and Pielak, 2017; Hesgrove and Boothby, 2020; Piszkiewicz et al., 2019). A model CAHS protein, CAHS D, is required for anhydrobiosis and provides desiccation protection when heterologously expressed in yeast and bacteria (Boothby et al., 2017). *In vitro*, CAHS D can protect lactate dehydrogenase (LDH) from denaturation when subjected to desiccation and rehydration (Boothby et al., 2017; Boothby and Pielak, 2017). However, a holistic molecular understanding of how CAHS proteins confer desiccation tolerance remains unknown.

A general mechanism proposed to explain desiccation tolerance is the vitrification hypothesis (Crowe et al., 1998). This hypothesis hinges on slowed molecular motion reducing the frequency and speed of damaging processes, such as protein unfolding. In the early stages of drying, protection is proposed to occur through inducing high viscosity in the system, slowing diffusion and molecular motion. Once in the vitrified solid state, molecular motion is slowed so dramatically that biological processes are essentially stopped, preventing further degradation of the system. While it has been shown that tardigrades and their CAHS proteins form vitrified solids, vitrification is not mutually exclusive with other potential mechanisms of desiccation tolerance (Boothby, 2021; Boothby et al., 2017; Boothby and Pielak, 2017; Hengherr et al., 2009; Hesgrove and Boothby, 2020).

Here we set out to further understand the mechanism(s) that underlie desiccation protection of client proteins by CAHS D. We present evidence that CAHS D can undergo a sol-gel transition in a concentration- and temperature-reversible manner. To understand how gelation impacts desiccation tolerance, we combined rational sequence design with a suite of complementary structural and biophysical techniques. Unexpectedly, we found that the mechanisms underlying protein stabilization do not correlate with the inhibition of protein aggregation, revealing that protection against these two major forms of protein dysfunction are mechanistically distinct. We also find that CAHS D’s interactions with water, as measured through T_2_ relaxation, was a strong predictor of unfolding protection. Our findings shed light not only on the fundamental biology underlying tardigrade anhydrobiosis and the function of IDPs during desiccation, but also provides avenues for pursuing applications such as the engineering of stress tolerant crops, and the stabilization of temperature sensitive therapeutics in a dry state.

## Results

### CAHS D undergoes a sol-gel transition

CAHS D (Uniprot: P0CU50) is a highly charged 227-residue protein that is predicted to be fully disordered (Boothby et al., 2017; Hesgrove and Boothby, 2020; Yamaguchi et al., 2012). During its expression and purification, it was observed that CAHS D undergoes a sol-gel phase transition, transitioning from a liquid into a solid gel state (Figure 1a). CAHS D gelation is concentration dependent: solutions below ∼10 g/L (0.4 mM) remain diffuse, solutions 10 g/L − 15 g/L are increasingly viscous, and above ∼15 g/L (0.6 mM), form robust gels (Figure 1a).

**Figure 1:**
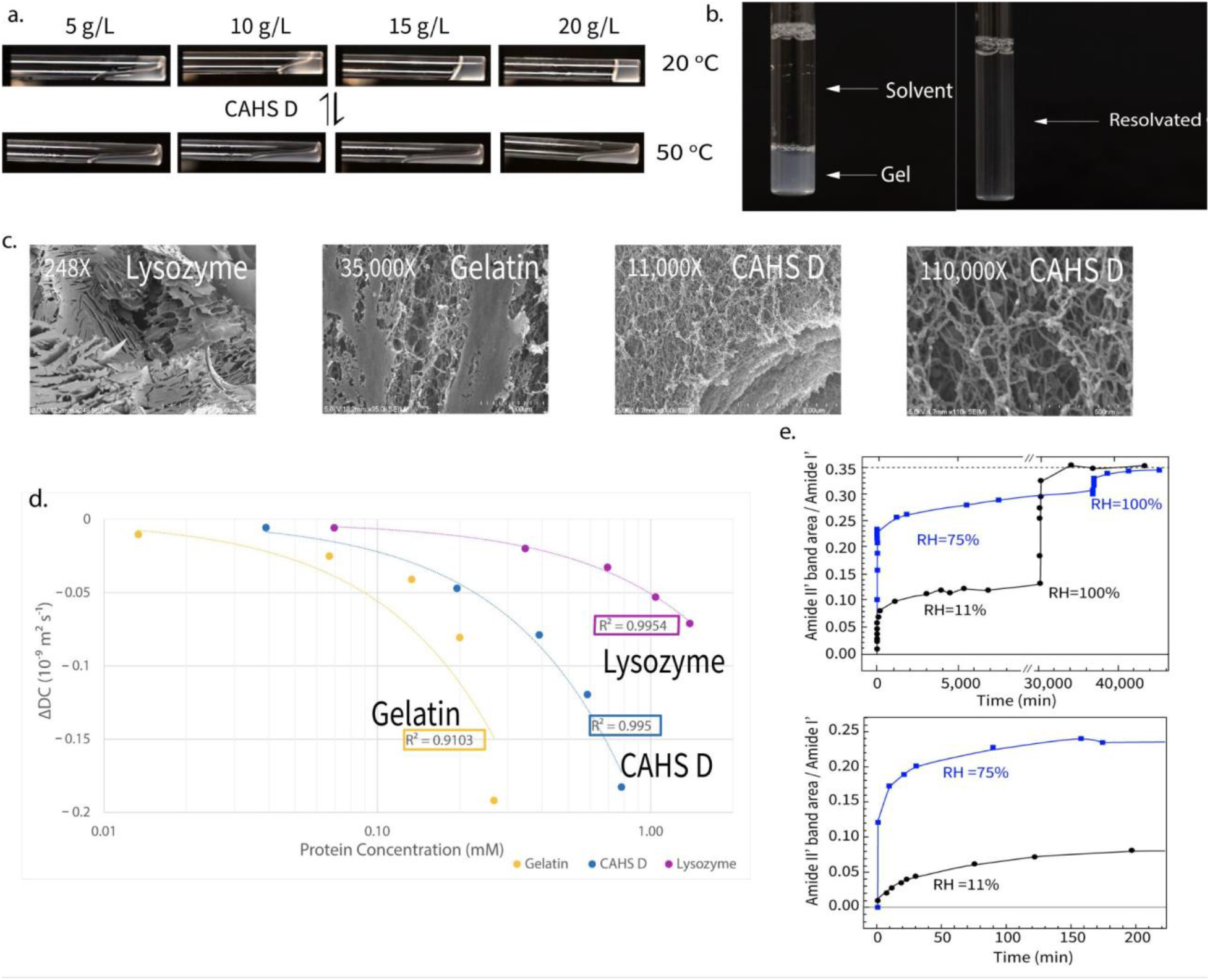
Gel properties of CAHS D. a) Reversible temperature and concentration dependent gelation of CAHS D. CAHS D at 20°C (top panel) shows gelation beginning at 15 g/L (0.6 mM), while at 50°C gelation is not observed in any concentration. Once cooled, gel formation recurs at 15 g/L. b) Dilution of 20 g/L (0.8 mM) CAHS D gel in 20mM Tris buffer results in resolvation. c) SEM images of lysozyme (248X), gelatin (35,000X), and CAHS D (11,000X and 110,000X). Images show the reticular nature of CAHS D gel structure is similar to that of gelatin. All SEM imaging was performed with proteins at 50 g/L. d) Δ Diffusion coefficients for lysozyme (pink), CAHS D (blue), and gelatin (gold). Gelled proteins show more dramatic slowing of ΔDC than non-gelling lysozyme, in a concentration dependent manner. Linear regressions for the full concentration range are shown as dotted lines, with R^2^ presented for each fit. A minimum of three measurements were used to calculate average diffusion values for all proteins at all concentrations. e) Kinetics of amide HDX in CAHS D glassy matrices. Left panel: CAHS vitrified gels at uniform initial hydration level were exposed to a D_2_O atmosphere at RH=11% (black symbols) or RH=75% (blue symbols). Subsequently both were transitioned to pure D_2_O atmosphere (RH=100%), which demonstrated that both dried gels showed similar HDX when fully saturated (supplemental text for more details). The dashed line represents the value of the amide II’ band area normalized to the area of the amide I’ band area. Right panel is an expansion of the left panel, showing the initial phase of the HDX kinetics. Source data for Figure 1 is available in files: Figure 1d - Source Data 1.xlsx and Figure 1e - Source Data 1.xlsx.

Gelation is reversible through heating (Figure 1a) and dilution (Figure 1b). Reversibility suggests that gelation is driven by non-covalent physical crosslinks, as opposed to chemical crosslinks which would yield an irreversible gel (Almdal et al., 1993). The thermal dependence of the sol-gel transition suggests that favorable enthalpy drives gelation, rather than the hydrophobic effect, since hydrophobic interactions are stabilized by increasing temperature (Dignon et al., 2019; Dill et al., 1989; van Dijk et al., 2015). This is reinforced by the reversibility seen through resolvation (Figure 1b), which would not lead to dissolution of hydrophobic interactions. Thus, gelation is driven by non-hydrophobic, non-covalent interactions such as hydrogen bonding, polar interactions, or charge interactions (Dignon et al., 2019).

High-resolution imaging reveals that CAHS D gels form reticular networks (Figure 1c). A fine meshwork of CAHS D fibers is interspersed with large pores (Figure 1c, Figure 1 – figure supplement 1a bottom left). This topology is similar to gels formed by gelatin, (Figure 1c) and morphologically distinct from crystalline solids formed by lysozyme (Figure 1c). The macromolecular architecture of CAHS D gels is reminiscent of that formed by synthetic polymers, in which relatively sparse physical crosslinks underlie the network connectivity (McComb et al., 2019). This implies that specific regions of CAHS D may drive intermolecular interactions.

Further investigation of the reticular mesh of the gel formed by CAHS D was pursued with small-angle X-ray scattering (SAXS) using a concentration gradient of the protein (Figure 1 - figure supplement 1a). We observed the concentration-dependent emergence of a scattering peak, indicative of a repeating structure approximately 9.5 nm in size (Figure 1 - figure supplement 1a top panels). This feature size corresponds well with the ∼10 nm fibers observed in our SEM imaging (Figure 1 - figure supplement 1a top-left panel). SAXS also allowed us to determine the mesh-size within the fibers, which shrunk in a concentration-dependent manner from ∼26 Å at low CAHS D concentration, to ∼20 Å at higher concentrations, consistent with greater structural integrity at higher concentrations (Figure 1 - figure supplement 1a bottom-right panel).

### Gelation Slows Diffusion, Vitrification Immobilizes Proteins

Slowed molecular motion through high viscosity and eventual vitrification is a cornerstone hypothesis in desiccation tolerance (Buitink and Leprince, 2004; Crowe et al., 1998). To determine how molecular motion is impacted in CAHS D gels, we assessed the diffusion of water in solutions of varying CAHS D concentration using low-field time-domain nuclear magnetic resonance spectroscopy (TD-NMR). This method is commonly used to determine the diffusion of water and oils within liquid, gel, and even solid materials (Alacik Develioglu et al., 2020; Rondeau-Mouro et al., 2019).

CAHS D dramatically slowed the diffusion of water, both below and above its gelation point (15 g/L, 0.6 mM) (Figure 1d). Gelatin, which forms gels through entwined triple helices, slowed diffusion more than CAHS D. In contrast, lysozyme, a non-gelling protein, slowed diffusion much less than both gelling proteins (Figure 1d).

To probe the degree to which hydration influences immobilization of the vitrified gel matrix, we performed hydrogen deuterium exchange (HDX) on dried and rehydrated gels at low and high relative humidities (RH=11% and RH=75%, respectively; Figure 1e). HDX experiments can distinguish between a tightly packed, conformationally restricted matrix (slow exchange), and a loosely packed pliable matrix (rapid exchange). At both relative humidities, the kinetics appear biphasic (Figure 1e). The major phase occurs quickly for both samples, but much faster in RH=75% (minutes) than RH=11% (tens of minutes) (Figure 1e bottom panel). The minor phase, present in both samples, is much slower (days). The limited total exchange (37%) observed at RH=11% is likely due to a substantial fraction of the CAHS amide groups in the matrix being excluded from contact with the deuterated atmosphere. Such an exclusion indicates tight packing of the CAHS protein within the matrix, and strong inhibition of conformational fluctuations decreasing accessibility of peptide amide groups to the deuterated atmosphere. This is consistent with the observation that, when incubated at RH=75%, the fraction of deuterated amide groups (83%) is more than doubled. The higher hydration level of the matrix at RH=75% (see the stretching D_2_O band in Figure 3 – figure supplement 1c) plasticizes the protein, increasing conformational fluctuations and amide accessibility to atmosphere. When both samples are kept at RH=100%, they both achieve the same level of HDX as a solubilized protein (Figure 1e, dashed line), indicating that the total amides competent for exchange in each protein sample is equivalent when fully hydrated. Therefore we conclude that increased HDX rates of CAHS D amides (Figure 1e) were strongly correlated with humidity, demonstrating dehydration-dependent immobilization of the CAHS D vitrified matrix.

In summary, biochemical, imaging, and biophysical assessment of CAHS D demonstrate that this protective protein undergoes a sol-gel transition, likely driven by assembly through non-covalent, non-hydrophobic interactions, and that the dynamics of the gel and embedded client proteins is dramatically influenced by the level of residual moisture in the system. The strong responsiveness of CAHS D to hydration and its ability to slow water diffusion has implications for the molecular mechanisms of client protein protection.

### Dumbbell-like Ensemble of CAHS D

CAHS proteins are highly disordered (Boothby et al., 2017; Hesgrove and Boothby, 2020), so standard structural studies are not feasible (Boothby et al., 2017; Hesgrove and Boothby, 2020). Instead, we performed all-atom Monte Carlo simulations to assess the predicted ensemble-state adopted by monomeric CAHS D proteins. Simulations revealed a dumbbell-like ensemble, with the N- and C-termini of the protein forming relatively collapsed regions that are held apart from one another by an extended and highly charged linker region (LR) (Figure 2a, Movie 1). Moreover, meta-stable transient helices are observed throughout the linker region (LR), while transient β sheets are observed in the N- and C-terminal regions (Figure 2a, Figure 2 – figure supplement 1a&b).

**Figure 2.**
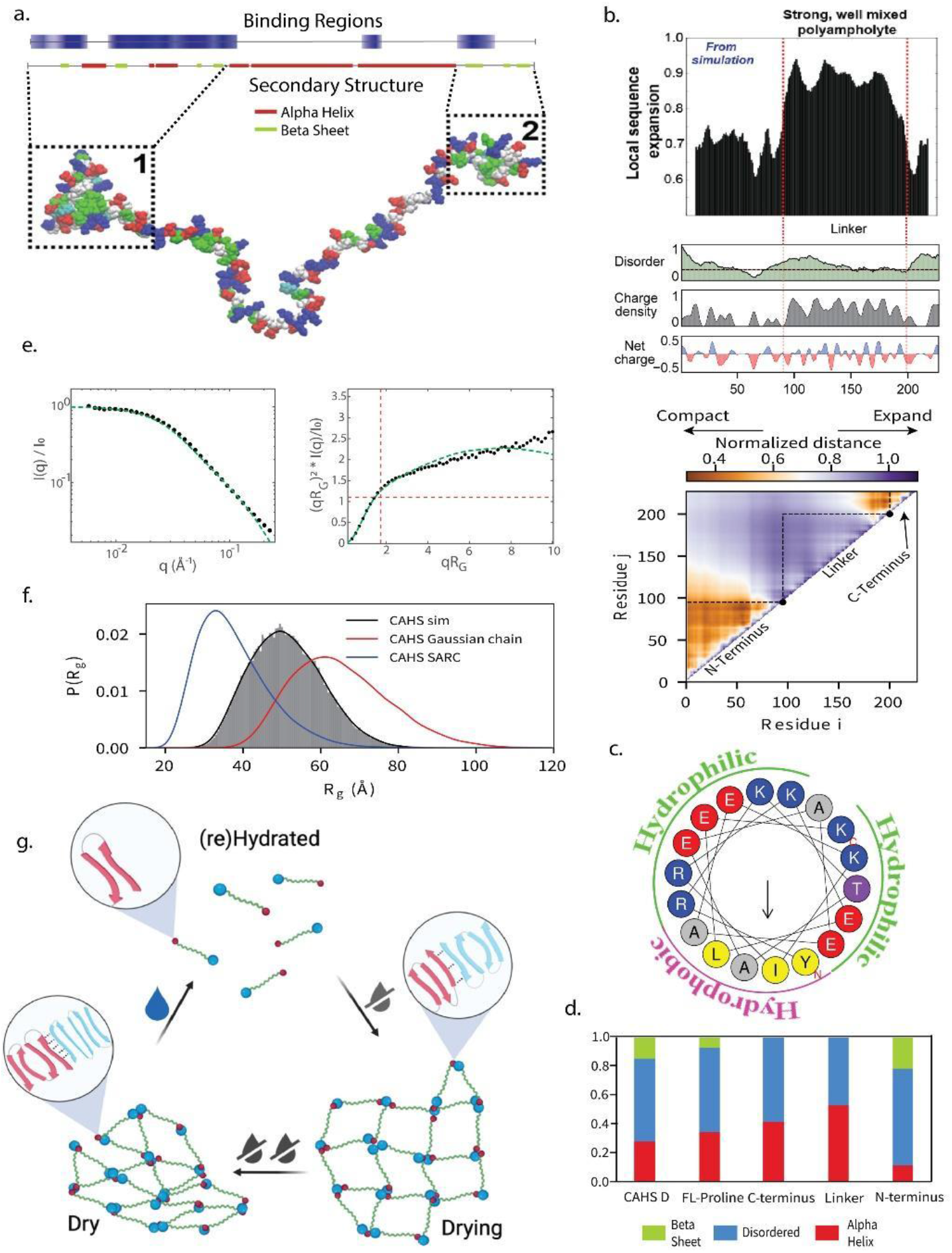
CAHS D gelation is driven by the stickers and spacers model. a) Bioinformatic predictions of secondary structure characteristics (top) and representative conformational global ensemble model (bottom) of CAHS D showing the extended central linker and sticky termini. b) Top panels illustrate the relationship between sequence expansion, disorder, charge density, and net charge of CAHS D. Heat map (bottom) shows normalized distance between residues as predicted in simulations. c) Helical wheel plot (HeliQuest, plothelix) shows the distribution of charged faces in a predicted linker helix. Colors represent the amino acid type (yellow, hydrophobic; grey, non-polar; red, acidic; and blue, basic). The arrow indicates the direction of the hydrophobic moment. d) The ratio of α helix, disorder, and β sheet secondary structure propensity determined by CD spectroscopy for CAHS D, FL-Proline, C-terminus, Linker and N-terminus. e) Raw SAXS data for monomeric CAHS D protein (left) and the Kratky transformation (right). Experimental data (black circles) and the form factor calculated from simulations (green lines) were normalized to the zero-angle scattering and overlaid. f) CAHS D (black) radius of gyration determined from simulations, compared radii of a self-avoiding random coil (blue) and a gaussian chain (red), of equal linear size. g) Proposed mechanism of gelation for WT CAHS D. As water is removed from the system, monomers assemble through β-β interactions in the termini. These interactions are strengthened as drying progresses. Upon rehydration, CAHS D gel can easily disassemble as seen in Figure 1c. Source data for Figure 2 is available in files: Figure 2d - Source Data 1.xlsx, Figure 2e – Source Data 1.zip and on GitHub.

To validate simulations, we performed SAXS on monomeric CAHS D. The radius of gyration (*R*_*g*_) – a measure of global protein dimensions – we obtained was 4.84 nm (simulation R_g_ = 5.1 nm), and the aligned scattering profiles obtained from simulation and experiment show good agreement (Figure 2e, Figure 2 – figure supplement 1c). These dimensions are substantially larger than those expected if a protein of this size was either folded (*R*_*g*_ = 1.5 - 2.5 nm) or behaved as a Gaussian chain (*R*_*g*_ = 3.8 nm), yet smaller than a self-avoiding random coil (*R*_*g*_ = 6.5 nm) (Figure 2f).

The expanded nature of CAHS D derives from the LR, which contains a high density of well-mixed oppositely charged residues (Figure 2b, Figure 2 – figure supplement 1d), preventing its conformational collapse (Das and Pappu, 2013; Holehouse et al., 2017). The transient helices formed in the LR have an amphipathic nature, (Figure 2a-c, Figure 2 – figure supplement 1a) and are predicted to have a hydrophobic and a charged face (Hesgrove and Boothby, 2020) (Figure 2c). We note that the CAHS D linker is among the most well-mixed, high-charge sequences in the entire tardigrade proteome (Figure 2 – figure supplement 2c); thus the extended nature of this sequence likely represents a functional, evolutionarily selected trait.

Circular dichroism (CD) spectroscopy confirms the largely disordered nature of full-length CAHS, with some propensity for α-helical and β-sheet formation (Figure 2d). CD spectroscopy performed on truncation mutants containing only the LR or N-terminal region confirmed substantial helical content in the LR (∼50%), and β-sheet content in the N-terminal region (∼20%) (Figure 2d). We observed no residual structure by CD in the isolated C-terminal region, contrary to the predicted β-sheet content. This could be caused by the loss of sequence context in the truncated terminus, so we inserted three structurally disruptive prolines (Imai and Mitaku, 2005; Williams et al., 2004) into the predicted C-terminal β-sheets of the full-length protein (Figure 3 – figure supplement 1a). CD on FL-Proline showed a ∼50% reduction in β-sheet content relative to wildtype (Figure 2d; Figure 1 – figure supplement 1b), confirming the β-sheet nature of the C-terminus.

Overall these data indicate that CAHS D exists in a dumbbell-like ensemble, which moves through conformational states consisting largely of β-sheeted termini held apart by an extended α-helical linker.

### Gelation is Driven by Terminal Interactions

A dumbbell-like protein with beta-sheeted termini would be an ideal candidate for gelation via the ‘stickers and spacers’ model (Choi et al., 2020; Li et al., 2012). In this model, discrete sites along a protein that contribute attractive intermolecular interactions are designated as stickers while non-interactive regions are spacers. Here, the N- and C-terminal regions of CAHS D can be considered stickers, while the LR is a spacer. When the spacer has a large effective solvation volume, like the expanded linker of CAHS D, phase separation is suppressed in favor of a sol-gel transition (Harmon et al., 2017). Moreover, we predict that intra-protein terminal interactions, which would suppress assembly through valence capping, are reduced by the separation enforced through the extended LR (Figure 2c) (Sanders et al., 2020). Based on our biophysical characterization of the monomeric protein, we hypothesized that gel formation of CAHS D occurs through inter-protein β-β interactions mediated between termini (Figure 2g).

To test this, we generated a range of CAHS D variants (Figure 3a) which disrupt the dumbbell-like ensemble, and thus should not gel. Consistent with our hypothesis that β-β interactions drive gelation, all variants lacking at least one termini resulted in a loss of gelation (N, LR, FL-Proline, NL1, CL1; Figure 3a, Figure 3 – figure supplements 3a&b). Unexpectedly, variants that replaced one terminal region for another (NLN and CLC; Figure 3a) also did not form gels under the conditions tested. These results show that heterotypic interactions between N- and C- termini are crucial for strong gel formation, implicating molecular recognition and specificity encoded by the termini. Interestingly, the 2X Linker variant, which maintained heterotypic termini but doubled the length of the linker, gelled rapidly at 5 g/L (0.1 mM), well below the gelling point of the wildtype protein at 15 g/L (0.6 mM) (Figure 3a). Thus, the length of the linker may tune the gel point by determining the monomeric molecular volume, setting the overlap concentration, which is a key determinant of the gel point (Colby, 2010; Sakumichi et al., 2021).

**Figure 3:**
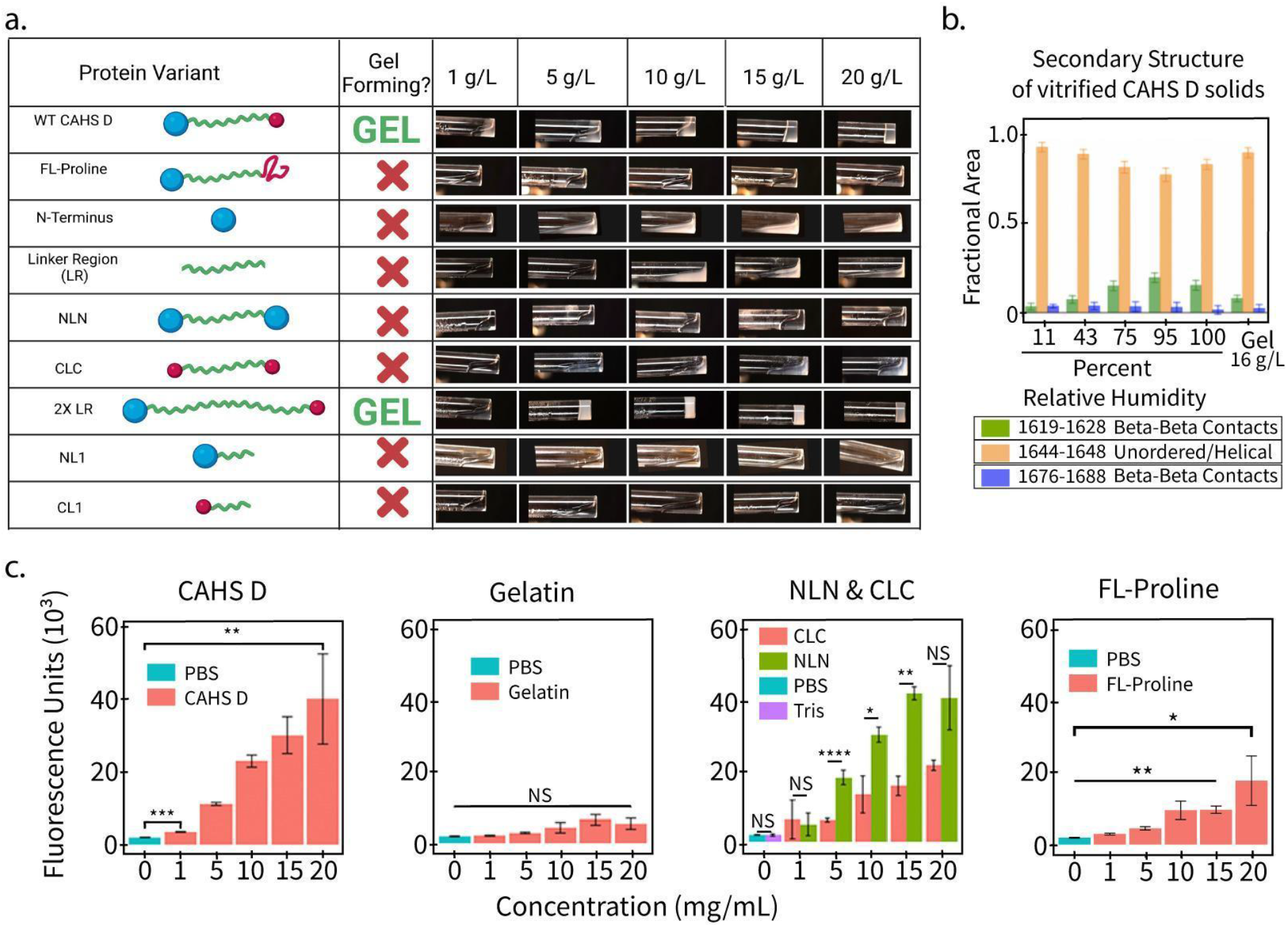
Gelation of CAHS D relies on its dumbbell-like ensemble, and intermolecular β-β interactions. a) Schematic of the variant structure, the gel propensity of each listed concentration, and photos of the proteins in solution at indicated concentrations. Only CAHS D and the 2X Linker variant showed gelation. Future depictions of these constructs include a green “G” next to the cartoon to indicate their ability to form a gel. b) Relative secondary structure content of CAHS D as determined by FTIR analysis of the amide I’ band in a glassy matrix at different hydration levels, and in the hydrated gel state at a concentration of 16 g/L (0.6 mM). The Gaussian sub-bands centered in the wavenumber interval (1644-1648 cm^-1^) are attributed to unordered/helical regions, while the Gaussian components peaking in the wavenumber intervals (1619-1628 cm^-1^) and (1676-1688 cm^-1^) are indicative of interprotein β-sheet structures. c) Thioflavin T (ThT) fluorescence as a function of concentration for CAHS D, gelatin, NLN, CLC and FL-Proline. Error bars represent standard deviation. Significance determined using a Welch’s t-test. All experiments presented used a minimum of 3 replicates. Asterisks represent significance relative to wild type CAHS D. *p<0.05, **p<0.01, ***p<0.001, ****p<0.0001, NS is not significant. Source data for Figure 3 is available in files: Figure 3b - Source Data 1.xlsx and Figure 3c - Source Data 1.xlsx.

### Optimal Hydration for β-β Stability

Using Fourier-transform infrared spectroscopy (FTIR) we observed that β-sheet interactions are maximal in CAHS glasses at 95% relative humidity (RH), and lowest in fully hydrated gels and glasses at 11% RH (Figure 3b). This raises the possibility that there is an optimal hydration level for stabilizing β-β contacts, which may relate to the need for higher stability while the matrix is undergoing drying or rehydration.

To confirm the role of β-β interactions in gelation, we assayed CAHS D solutions at ranging concentrations with thioflavin T (ThT). ThT is used as a fluorescent indicator of amyloid fibrils (Biancalana et al., 2009; Wu et al., 2009), and can report on β-β interactions (Ge et al., 2018; Namioka et al., 2020; Peccati et al., 2017). We observed increases in ThT fluorescence intensity as a function of CAHS D concentration, with the most dramatic increase between 5 - 10 g/L (0.2 - 0.4 mM), suggesting nucleation & assembly of CAHS D monomers prior to gelation at 15 g/L (0.6 mM) (Figure 3c). Gelatin did not show concentration dependent changes in ThT fluorescence and was not significantly different from buffer controls (Figure 3c). ThT labeling of NLN and CLC variants showed a concentration-dependent increase in fluorescence, with more interactions shown in NLN than CLC (Figure 3c), consistent with the predicted degree of β-sheet content in each (Figure 2a&d). FL-Proline β-β interactions were lesser than variants with two folded termini, as expected. These data suggest that β-β interactions increase parallel to gelation and that assembly events are occurring between monomers, prior to an observed system-wide sol-gel transition.

Together, our SAXS data and bioinformatic analyses show that the extended nature of the protein can be attributed to the LR, and that this extension helps to hold the termini of an individual protein apart. CD, FTIR, and ThT labeling data show that CAHS D gelation is mediated by inter-protein β-β contacts formed through the interaction of the termini, which are influenced by hydration.

### Mechanistic Determinants of Protection

Protein gelation is uncommon in a biologically relevant context, and in these instances is often functional (Böni et al., 2017; Kesimer et al., 2010). This led us to wonder if gelation of CAHS D is linked to protection. The enzyme lactate dehydrogenase (LDH) unfolds and becomes irreversibly non-functional (Boothby et al., 2017; Piszkiewicz et al., 2019; Rani, 2019), but does not aggregate (Chakrabortee et al., 2012; Popova et al., 2015), during desiccation and an LDH activity assay is commonly employed to measure protection against unfolding (Boothby et al., 2017; Piszkiewicz et al., 2019; Popova et al., 2015).

In contrast, citrate synthase (CS) forms non-functional aggregates after successive rounds of dehydration and rehydration (Chakrabortee et al., 2012). We tested all gelling and non-gelling variants (Figure 3a) using both assays, to determine how gelation influences CAHS D protective capacity (Figure 4a&b, Figure 4 – figure supplement 1), and were surprised to see that while gelling variants prevented unfolding best, they performed least well at preventing aggregation (Figure 4a&b, Figure 4 – figure supplement 1), suggesting that different forms of protein dysfunction are prevented through distinct mechanisms.

**Figure 4:**
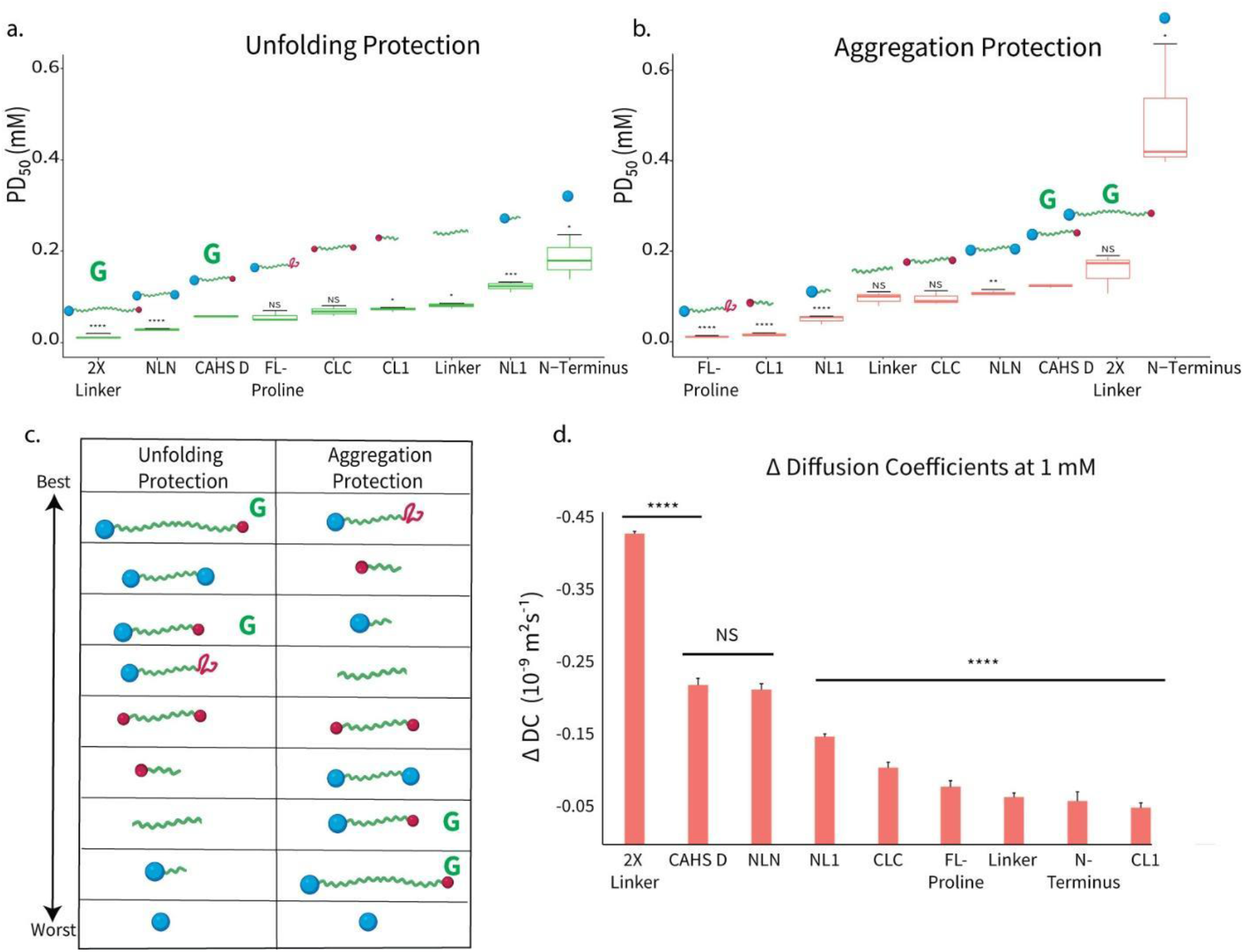
Gelation promotes protection against protein unfolding, but not aggregation, during desiccation. Unfolding(a) and aggregation (b) PD50 values (shown left to right, in order from best to worst) of each variant.. Lower PD50s correspond with better protective capability. Significance was determined for both (a) and (b) using a Welch’s t-test, of variant’s protection relative to that of wildtype. c) Ranking of variants in both the unfolding and aggregation assays, with the lowest PD50 (best protectant) at the top and the highest PD50 (worst protectant) at the bottom. Refer to (a) and (b) for information on relative protective ranking of each variant. Green G indicates that a variant is gel forming. d) ΔDiffusion Coefficients for all variants at 1 mM, calculated from linear fits for each variant’s full concentration range (1-20 mg/mL). Error bars represent the standard error for the full concentration range for each variant. Significance was determined as a *χ*^2^ analysis of the equality of linear regression coefficients for the linear fit of each variant, compared to that of CAHS D. All experiments used a minimum of 3 replicates. Asterisks represent significance relative to the wildtype.*p<0.05, **p<0.01, ***p<0.001, ****p<0.0001, NS is not significant. Source data for Figure 4 is available in files: Figure 4a - Source Data 1.xlsx, Figure 4b - Source Data 1.xlsx and Figure 4d - Source Data 1.xlsx.

### Determinants of Aggregation Protection

The mechanism most often attributed to the prevention of desiccation-induced protein aggregation is the molecular shielding hypothesis (Chakrabortee et al., 2012; Furuki et al., 2020; Hatanaka et al., 2013). This posits that protein aggregation can be prevented by protectants that act as disordered spacers, which impede interactions between aggregation prone molecules (Chakrabortee et al., 2012; Furuki et al., 2020; Goyal et al., 2005; Hatanaka et al., 2013). Shielding proteins generally have some means of interacting with client proteins, although the interactions are weak (Chakrabortee et al., 2012; Furuki et al., 2020; Hatanaka et al., 2013; Ikeda et al., 2020).

Results from our CS aggregation assay are consistent with this mechanism; the presence of extended LR with a single terminus emerged as the primary determinant of CS aggregation prevention (Figure 4b). Meanwhile, the presence of two folded termini seemed antagonistic to aggregation protection, suggesting that higher order assembly, made possible by two termini, is detrimental to preventing aggregation.

To determine the role of diffusion in protection, we measured diffusion of water and compared this with aggregation protection (Figure 4c,d). We found that neither aggregation protection nor diffusion were dependent on molecular weight or gelation. For example, FL-Proline (∼25 kDa) is best at aggregation protection, does not form gels, and has much faster water diffusion than CAHS D (∼25 kDa). On the other hand, NLN (∼31 kDa) slowed diffusion nearly identically to CAHS D, yet did not form gels, and ranked 6th at protecting from aggregation (CAHS D was ranked 7th) (Figure 4c,d).

We found that diffusion at 1mM normalized to the variant’s molecular weight loosely followed an inverse trend, where faster diffusion trended with lower aggregation PD50s (Figure 5d, Figure 5 – figure supplement 2d). In general, the higher linker content a variant contained, the better it performed at aggregation protection, so long as the variant also had a single folded terminus. Variants with a greater than 75% proportion of N-terminal content (NL1 and N-terminus), were outliers to this trend, both by having a high PD50 with fast diffusion (N-terminus), and by having a very low PD50 with slow diffusion (NL1). This is an interesting result that indicates that the N-terminus may in some way impact the relationship between diffusion and aggregation protection.

**Figure 5:**
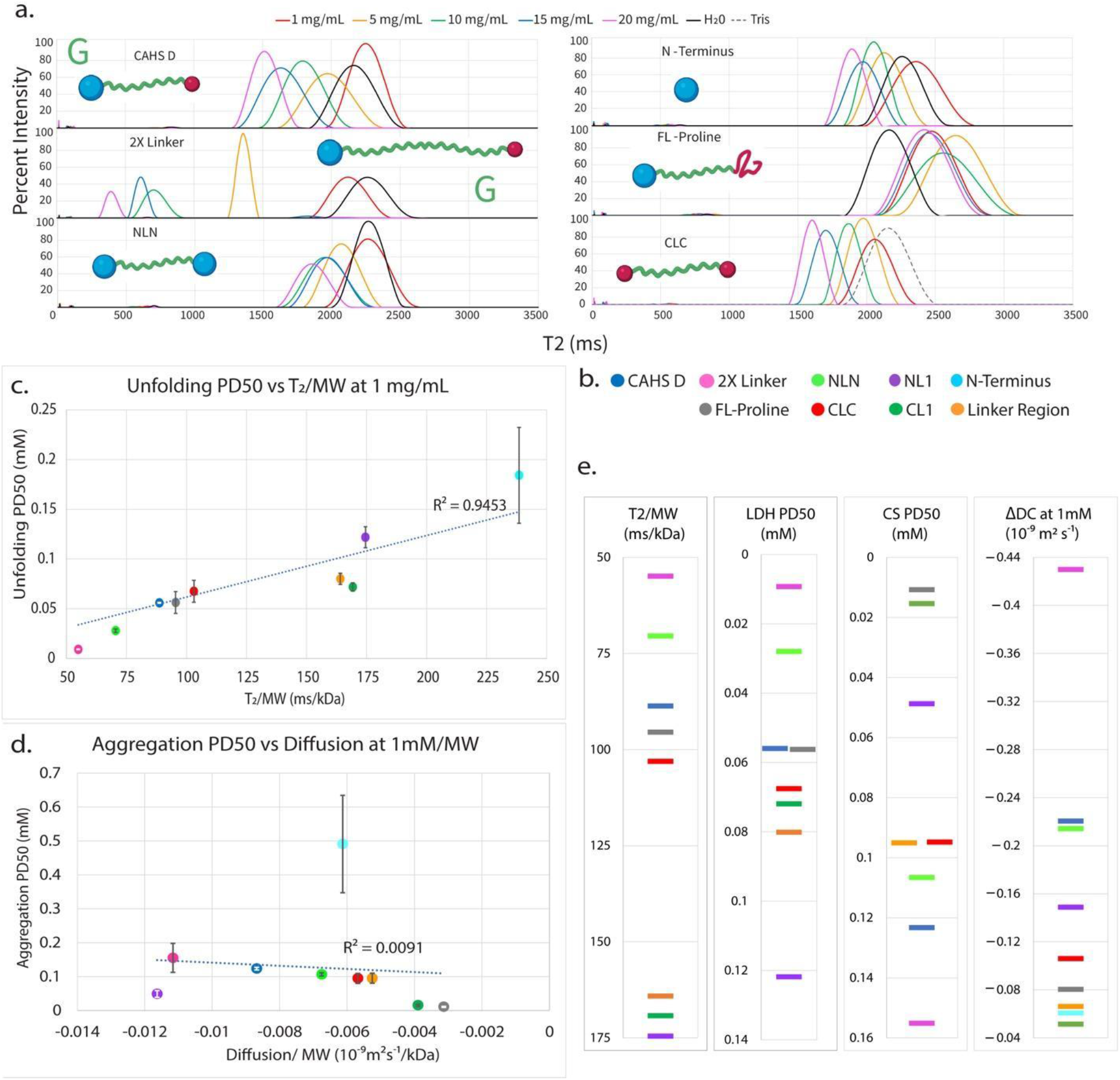
Slowed diffusion and coordination of water underlie CAHS D’s ability to promote protection against protein unfolding. a) T_2_ distributions of variants WT CAHS D, 2X Linker, N-terminus, NLN, LR, and CLC. (All variants not shown here can be found in Figure 5 – figure supplement 1b). 2X linker, CAHS D, and NLN show the most shifted T_2_ peak midpoints as a function of concentration. b) Legend for plots in (c-e). c) Unfolding PD50 as a function of water coordination per kilodalton for the 1 g/L. As shown by the linear trend of these plots, the water coordination per kilodalton is a strong indicator of unfolding protection. The relationship is shown in further detail in (e). d) Aggregation PD50 as a function of water diffusion at 1 mM per kilodalton. As shown, there is not a strong correlation between these properties. e) Single variable plots of Unfolding PD50, water coordination per kilodalton, Aggregation PD50, and ΔDC at 1mM. The top three variants that protect best against unfolding also show the slowest diffusion, however the relationship between ΔDC and unfolding protection breaks down after this point. Comparison of the Unfolding PD50 ranking and the T_2_/kDa ranking shows a striking relationship. However, no such relationship is seen for aggregation protection (Figure 5 – figure supplement 2a&d). Source data for Figure 5 is available in files: Figure 5a - Source Data 1.xlsx, Figure 5c - Source Data 1.xlsx, Figure 5d - Source Data 1.xlsx, and Figure 5e - Source Data 1.xlsx.

**Figure 6:**
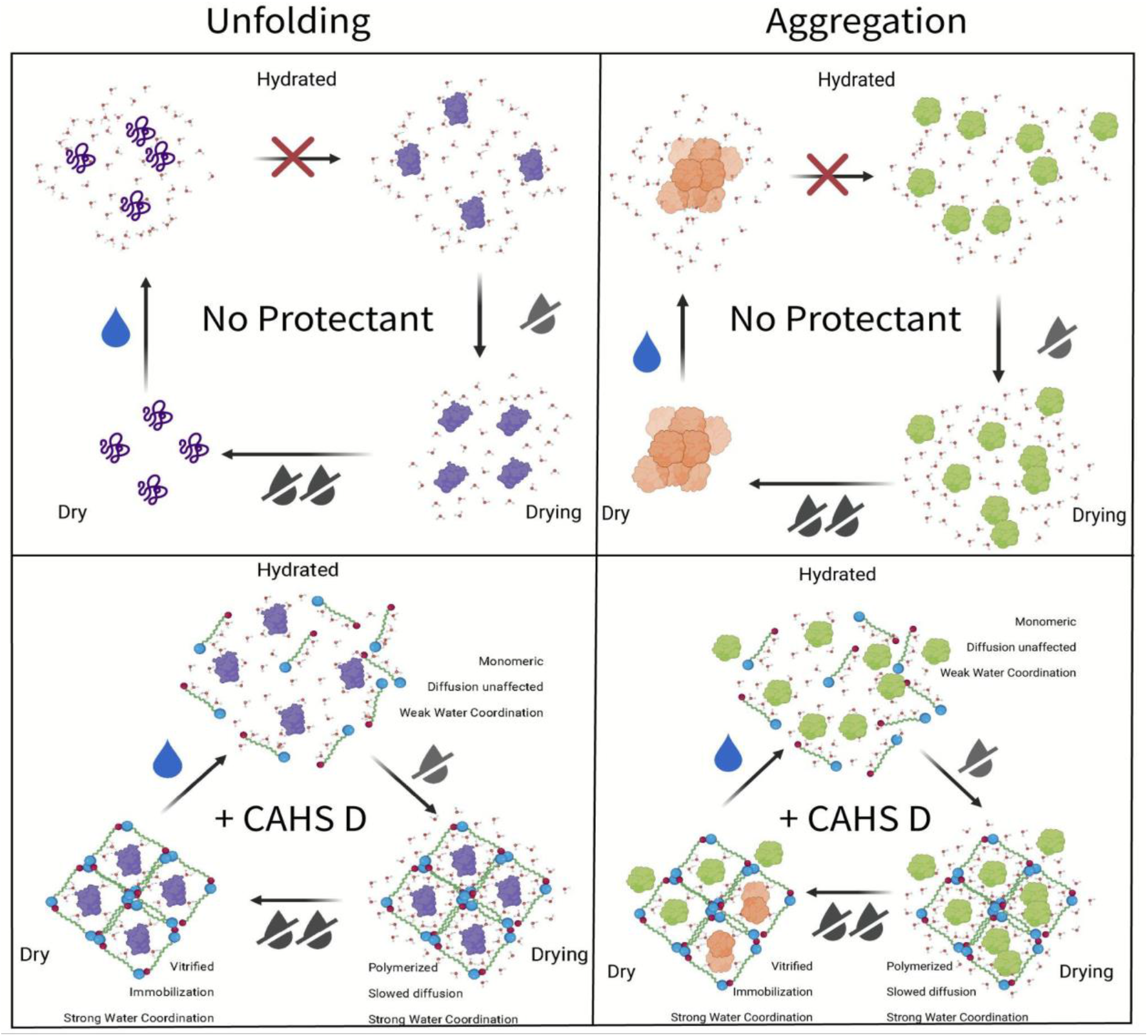
Proposed mechanism of protection against desiccation induced protein aggregation and unfolding. Schematics depict the proposed mechanisms occurring within the meshwork of CAHS D fibers. The left panels show what occurs in the case of protein unfolding, while the right panels illustrate protein aggregation. Top panels display the system without protectants, while the lower panel of each shows the system with wildtype CAHS D. As water is lost, self-assembly begins in samples with CAHS D. Water is coordinated along the linker and to some degree the N- terminus of CAHS D. As samples enter a vitrified state, some water remains coordinated by the linker, but most water has been lost. Upon rehydration, the matrix readily disassembles and releases molecules from the gel structure. Inferred behaviors of CAHS variants can be found in Figure 6 – figure supplement 1 and Figure 6 – figure supplement 2 for protein unfolding and aggregation, respectively.

The negative relationship between diffusion and aggregation protection conflicts with key aspects of the vitrification hypothesis. However, if we consider aggregation protection to be primarily driven through molecular shielding, then this antagonism between anti-aggregation activity and diffusion can be understood more clearly. Slowed diffusion would dampen dynamic movement of protectants around clients, limiting client isolation, making the protectant a less effective aggregation shield. This suggests that rapid diffusion may be a key part of molecular shielding.

### Determinants of Unfolding Protection

Gelation and diffusion also cannot fully explain protection against unfolding. For example, NLN did not gel, had diffusion identical to that of CAHS D, and was more protective against unfolding damage than CAHS D. Thus, neither diffusion nor gelation alone can predict unfolding protection.

A potential driver of unfolding protection is the amount of extended LR present in a variant. This is true both in dual-termini variants, and in single-termini variants. An example is found by the higher protection afforded by CL1 (9.7 kDa Linker content) versus NL1 (2.9 kDa Linker content). Monomer assembly, even if it does not ultimately lead to gelation, is critical for unfolding protection. All variants with two folded termini were either better or equivalent to CAHS D unfolding protection. This property, however, cannot predict protection displayed by variants with only one folded terminus.

Taken together, our LDH and CS assays show that linker content is a driver of protection in both forms of protein dysfunction. However, the context of the linker ultimately determines its functionality. The properties that drive unfolding protection are distinct from those that dictate aggregation protection (Figure 4c), again implying that the mechanisms underlying prevention of these unique forms of protein dysfunction are distinct.

### Water coordination predicts unfolding, but not aggregation, protection

Since gelation, assembly, linker content, and diffusion do not fully account for the protective trends observed in our unfolding assay, we looked to other mechanistic hypotheses regarding desiccation protection to explain how this form of protection might be mediated.

The water entrapment hypothesis addresses how a protectant may compensate for the loss of a stabilizing hydrogen bond network (HBN) experienced by a client protein during desiccation stress. This hypothesis proposes that the protectant coordinates a layer of water between itself and the client protein, to maintain the client’s HBN (Belton and Gil, 1994).

To understand how CAHS D and its variants interact with water, TD-NMR was used to measure T_2_ relaxation (Figure 5a). T_2_ relaxation yields information regarding the coordination of water molecules in a system (Ghi et al., 2002; Käiväräinen et al., 1984; Qiao et al., 2019; Wang et al., 2018; Zheng et al., 2006). Strongly coordinated water molecules are structurally organized, have reduced motion overall and less freedom of motion, and slower exchange within hydration layers (Adhikari et al., 2020; Ahmed et al., 2014; Laage et al., 2017; Lerbret et al., 2012; Pattni et al., 2017; Raschke, 2006). Therefore, faster T_2_ relaxation times correspond with coordinated or structured water molecules, and slower relaxations indicate less coordinated water, such as is found in bulk liquids (Figure 5 – figure supplement 1a).

Among our CAHS D variants, we observed a remarkable diversity of water coordinating properties (Figure 5a, Figure 5 – figure supplement 1b). Concentration-dependent water coordination was found, in order of strength, in 2X linker, CAHS D, CLC, NLN, and N-terminus (Figure 5a). For a deeper understanding of the relationship between water coordination and protection, we plotted the PD50 for our LDH and CS assays as a function of the major T_2_ peak midpoints normalized to the variant’s MW (T_2_/MW) at 1 g/L for all variants (Figure 5b,c). This represents what is essentially water-coordinating ability per kDa of protein, and gives insight into the degree to which a protein can coordinate water when the overall size of each protein is taken into account. We were surprised to find a strong correlation between T_2_/kDa and unfolding protection (Figure 5c) which was not replicated in aggregation protection data (Figure 5 – figure supplement 2a).

This further reinforces the notion that CAHS D uses disparate mechanisms to protect from different forms of protein damage; molecular shielding for aggregation protection and vitrification and water coordination for protein unfolding. The relationship between T_2_ relaxation and unfolding protection signals the relative importance of water coordinating capability in protecting a client protein’s native fold, which in turn provides evidence for the importance of accounting for hydration when considering the proposed mechanisms for desiccation protection.

## Discussion

In this work we set out to understand the drivers of CAHS D desiccation protection. We show that CAHS D forms reversible physical gels, an observation explained by a model in which attractive interactions encoded by the N- and C-terminal regions facilitate intermolecular self-assembly. In contrast, the large central linker region acts as an extended, highly charged spacer, reducing intramolecular interactions and setting the gel-point based on the overall protein dimensions. From a functional perspective, neither gelation nor diffusion correlate directly with prevention of protein aggregation or unfolding. Protection from unfolding is influenced by structural features but can be directly related to a variant’s ability to coordinate water. Aggregation protection, on the other hand, seems to be favored in systems with more rapid diffusion that do not form higher order assemblies.

Based on our results, we propose that water entrapment, wherein the protectant traps a layer of hydrating water between itself and the client protein (Belton and Gil, 1994; Shimizu and Smith, 2004), is the major mechanism driving CAHS D-mediated unfolding protection. Conversely, molecular shielding provides the best explanation of the trends observed for aggregation protection. We propose that CAHS D functions as a molecular Swiss Army Knife, offering multiple protective capabilities though distinct mechanisms.

### Dynamic Functionality for the Dynamic Process of Drying

Surviving desiccation is not about surviving a singular stress, but rather surviving a continuum of inter-related stresses. During early drying, the need to prevent aggregation may outweigh the need to maintain hydration of most cellular proteins. At this early stage, monomeric or low-oligomeric CAHS D could be performing important shielding functionalities to mitigate a cascade of aggregation. Such an aggregation cascade has been implicated in proteostatic dysregulation (Klaips et al., 2018). Aggregation protection during early drying would help preserve functional proteins during a critical window when the cell must adapt to water loss.

As drying progresses and further water is lost, CAHS D concentration increases(Boothby et al., 2017) and its primary function may transition from aggregation prevention to protection against unfolding. As small oligomers of CAHS D grow and combine, clients could become trapped between assembled CAHS D. This would enable CAHS D to hold coordinated water to the surface of the entrapped clients, hydrating them and maintaining their hydrogen bonding networks. These smaller assemblies would slow diffusion in the system more and more as they condense to form the gel fibers observed in SEM (Figure 1c).

This fibrous network of gelled CAHS D could provide a bridge from the drying phase to the fully dried vitrified solid phase. Molecular motion in a CAHS D gel is slower than a liquid, but faster than a vitrified solid, and residual water can be coordinated and held to the surface of client proteins. This would mitigate the water loss experienced by client proteins, allowing them to retain their native fold for as long as possible. By the time dehydration overwhelms CAHS D’s capacity to hold water, a vitrified solid will have formed. At this point, even though the client’s native fold may no longer be stabilized by a hydrogen bond network; full unfolding *still* would not occur because of the degree to which molecular motion is reduced in the vitrified solid.

The final phase of surviving desiccation is to rehydrate and return to active biological functions. CAHS D gels easily solubilize after gelation, indicating that they can rehydrate, resolvate, and release entrapped, protected clients, likely an essential step in efficiently returning the rehydrating organism to activity.

Our work provides empirical evidence that multiple mechanisms contribute to desiccation tolerance, and that these mechanisms can be mediated by a single protectant molecule.

## Acknowledgements

This work was supported by DARPA award W911NF-19-2-0019 and an Institutional Development Award (IDeA) from NIH grant (P20GM103432) to T.C.B. NIH award R35GM137926 to S.S.; Financial support from MIUR of Italy (RFO2019) is gratefully acknowledged by M.M., F.F. and G.V. A fellowship to S.B. and this work were partially funded by Wyoming NASA EPSCoR, NASA Grant #80NSSC19M0061. This research used resources of the Advanced Photon Source, a U.S. Department of Energy (DOE) Office of Science User Facility operated for the DOE Office of Science by Argonne National Laboratory under Contract No. DE-AC02-06CH11357 and resources supported by grant 9 P41 GM103622 from the National Institute of General Medical Sciences of the National Institutes of Health. Use of the Pilatus 3 1M detector was provided by grant 1S10OD018090-01 from NIGMS. Drs. Gary J. Pielak and Samantha Piszkiewicz are acknowledged and thanked for their early discussions and efforts, which were made possible with support from NIH award R01GM127291 to G.J.P. Lorena Rebecchi (University of Modena and Reggio Emilia, UNIMORE, Italy) is thanked for stimulating discussions and valuable advice.

## Materials and Methods

### Cloning

All variants and wild type CAHS D were cloned into the pET28b expression vector using Gibson assembly methods. Primers were designed using the NEBuilder tool (New England Biolabs, Ipswitch, MA), inserts were synthesized as gBlocks and purchased from Integrated DNA Technologies (Integrated DNA Technologies, Coralville, IA).

### Protein Expression

Expression constructs were transformed in BL21 (DE3) E. coli (New England Biolabs) and plated on LB agar plates containing 50 μg/mL Kanamycin. At least 3 single colonies were chosen for each construct and tested for expression.

Large-scale expression was performed in 1 L LB/Kanamycin cultures, shaken at 37°C (Innova S44i, Eppendorf, Hamburg, Germany) until an OD_600_ of 0.6, at which point expression was induced using 1 mM IPTG. Protein expression continued for four hours, after which cells were collected at 4000 g at 4°C for 30 minutes. Cell pellets were resuspended in 10 mL of resuspension buffer (20 mM tris, pH 7.5, 30 μL protease inhibitor [Sigma Aldrich, St. Louis, MO). Pellets were stored at -80°C.

### Protein Purification

Purification largely follows the methods in Piszkiewicz *et al*., 2019. Bacterial pellets were thawed and heat lysis was performed. Pellets were boiled for five minutes and allowed to cool for 10 minutes. All insoluble components were removed via centrifugation at 5,000 g at 10°C for 30 minutes. The supernatant was sterile filtered with 0.45 μm and 0.22 μm syringe filters (Foxx Life Sciences, Salem, NH). The filtered lysate was diluted 1:2 in purification buffer UA (8 M Urea, 50 mM sodium acetate [Acros Organics, Carlsbad, CA], pH 4). The protein was then purified using a cation exchange HiPrep SP HP 16/10 (Cytiva, Marlborough, MA) on the AKTA Pure 25 L (Cytiva), controlled using the UNICORN 7 Workstation pure-BP-exp (Cytiva). Variants were eluted using a gradient of 0-50% UB (8 M Urea, 50 mM sodium acetate, and 1 M NaCl, pH 4), over 20 column volumes.

Fractions were assessed by SDS-PAGE and pooled for dialysis in 3.5 kDa MWCO dialysis tubing (SpectraPor 3 Dialysis Membrane, Sigma Aldrich). For all variants except CLC, pooled fractions were dialyzed at 25°C for four hours against 2 M urea, 20 mM sodium phosphate at pH 7.0, then transferred to 20 mM sodium phosphate at pH 7 overnight. This was followed by six rounds of 4 hours each in Milli-Q water (18.2 MΩcm). Dialyzed samples were quantified fluorometrically (Qubit4 Fluorometer, Invitrogen, Waltham, MA), aliquoted in the quantity needed for each assay, lyophilized (FreeZone 6, Labconco, Kansas City, MO) for 48 hours, then stored at -20°C until use. CLC was dialyzed in 2 M urea, 20 mM Tris at pH 7 for four hours, followed by 6 rounds of 4 hours each in 20 mM Tris pH 7. CLC samples were quantified using Qubit4 fluorometer as described, concentrated using amicon spin-concentrators (Sigma-Aldrich) to the desired concentration and used immediately.

### Visual gelation and heat/dilution gel resolubilization assay

Quantitated and lyophilized protein samples were transferred as powder into NMR tubes (Wilmad Lab Glass, Vineland, NJ)and resuspended in 500 μL of water to a final concentration of 5 g/L, 10 g/L, 15 g/L, and 20 g/L. Samples were left at room temp for 5 minutes to solubilize. If solubilization was not occurring (as determined visually), samples were moved to 55°C for intervals of 5 minutes until solubilized. If solubilization was not progressing at 55°C after 10 minutes of heating (as determined visually), then samples were transferred 95°C for 5-minute intervals until fully solubilized.

Solubilized proteins were transferred from heat blocks to the bench and left at ambient temperature for 1 hour. Tubes were then loaded horizontally into a clamp holder and photographed. Gelation was visually assessed by the degree of solidification or flow of the sample in the NMR tube.

After 1 hour at ambient temperature, proteins that had been found to form gels were transferred to a 55°C heat block for 1 hour. After an hour, samples were returned to the photographic clamp holder and imaged immediately to confirm that gelation had been disrupted. Sample where placed upright on the bench at ambient temperature for 1 hour and allowed to reform gels and then imaged again as above.

A duplicate sample of 20 g/L CAHS D was prepared as described and assayed for dilution resolubility. Buffer (Tris, 20 mM pH 7) was added to the gel to bring the final concentration of solvated CAHS D to 5 g/L, which is below the gelation point of the protein. Sample was photographed immediately after addition of buffer, vortexed for 5 seconds, and left to resolubilize. Sample completely dissolved within 30 minutes of buffer addition.

### Helical Wheel Generation

Helical wheel plots were generated using the heliQuest sequence analysis module. CAHS D linker sequence 126-144 was used, with the α-helix option chosen as the helix type to model.

### Scanning electron microscopy and critical point drying

Protein samples were heated to 95°C for 5 minutes and 50 μl of each sample were transferred to microscope coverslips. Samples were fixed in a 2.5% glutaraldehyde / 4% paraformaldehyde solution for 10 minutes. Samples were then dehydrated in an ethanol series going from 25%, 50%, 75%, 3x 100% with each incubation time being 10 minutes. Dehydrated samples were prepared for imaging using critical point drying (Tousimis Semidri PVT-3, SGC Equipment, Austin, TX) and supporter coating (108 Auto, Cressington Scientific Instruments, Watford, United Kingdom). Imaging was performed on a Hitachi S-47000 scanning electron microscope.

### Amide hydrogen/deuterium exchange kinetics in a CAHS glassy matrix at different hydration levels

Amide H/D exchange can be followed by using the amide II band of proteins, a mode which is essentially the combination of the NH in-plane bending and the CN stretching vibration. Following H/D exchange the N–D bending no longer couples with the CN stretching vibration converting the mode to a largely CN stretching vibration around 1450 cm^-1^ (Barth, 2007). The wavenumber of this mode (the so-called amide II’ band) is approximately 100 cm^-1^ downshifted compared to the amide II mode detectable in H_2_O (around 1550 cm^-1^). Thus, the two bands are clearly separated in the spectrum.

In order to examine the extent and kinetics of the amide H/D exchange we prepared two samples of lyophilized CAHS protein in H_2_O, then extensively dried the gel by equilibration within gas-tight sample holders at RH=6% in the presence of NaOH.H_2_O. The attainment of a minimum, steady hydration level was checked by monitoring (v_2_+v_3_) combination band of water around 5150 cm^-1^ (Malferrari et al., 2011) over 3 days, using the amide I band as an internal standard. After three days of equilibration, H/D exchange at RH=11% was begun by placing saturated LiCl in D_2_O in one of the CAHS D sample holders to create an 11% D_2_O atmosphere. A series of FTIR spectra was then recorded in sequence, at selected time intervals following the start of H/D exchange. For each spectrum, 100 interferograms were averaged to allow a sufficiently rapid acquisition (3 minutes) particularly at the beginning of the H/D exchange. This process was performed in parallel on the second sample, where H/D exchange at RH=75% was begun by placing a saturated solution of NaCl in D_2_O in the second sample holder.

Figure 3 – figure supplement 1e shows a series of spectra in the amide region (1350 – 1750 cm^-1^) acquired in the latter experiment at selected time intervals from the time (t=0) when H/D exchange started, following incubation in the presence of the saturated NaCl solution in D_2_O. We have previously shown (Malferrari et al., 2016; Malferrari et al., 2012) that such an isopiestic approach for isotopic exchange is quite effective and rapid (hour time scale).

As expected for a progressive amide H/D exchange, an intense amide II’ band centered around 1450 cm^-1^ appears within minutes. The progressive appearance and increase of the amide II’ band is accompanied by a progressive absorbance decrease in the spectral region of the amide II band at 1550 cm^-1^. This partially masks the appearance of a band peaking around 1575 cm^-1^, attributed to side chain vibrations, which is clearly detected in the fully deuterated glass (see Figure 3 – figure supplement 1c). This band has been assigned to an overlap of v_as_(COO^-^) from Asp and Glu and of Arg v_s_(CN_3_H_5_^+^) (Barth, 2007; Goormaghtigh et al., 2016). We evaluated the extent of amide H/D exchange from the area of the amide II’ band at 1450 cm^-1^ after subtraction of a straight baseline drawn between the minima on either side of the band (Goormaghtigh et al., 1994). The contribution of the small background band present at t=0 was also subtracted, and the resulting area was normalized to the area of the amide I/I’ band.

### Small-angle X-ray scattering

All SAXS measurements were performed at the BioCAT (beamline 18ID at the Advanced Photon Source, Chicago, IL). SAXS measurements on monomeric CAHS were collected with in-line size exclusion chromatography (SEC-SAXS) coupled to the X-ray sample chamber to ensure the protein was monomeric. Concentrated protein samples were injected into a Superdex 200 increase column (Cytiva) pre-equilibrated in a buffer containing 20 mM Tris pH 7, 2 mM DTT, and 50 mM NaCl. Scattering intensity was recorded using a Pilatus3 1 M (Dectris) detector placed 3.5 m from the sample, providing a q-range from 0.004-0.4 Å^-1^. One-second exposures were acquired every two seconds during the elution. Data were reduced at the beamline using the BioXTAS RAW 1.4.0 software (Hopkins et al., 2017). The contribution of the buffer to the X-ray scattering curve was determined by averaging frames from the SEC eluent, which contained baseline levels of integrated X-ray scattering, UV absorbance, and conductance. Baseline frames were collected immediately pre- and post-elution and averaged. Buffer subtraction, subsequent Guinier fits, and Kratky transformations were done using custom MATLAB (Mathworks, Portola Valley, CA) scripts.

CAHS samples were prepared for SAXS measurements by dissolving 5 mg/mL lyophilized CAHS protein into a buffer containing 20 mM Tris pH 7 and 50 mM NaCl. Samples were incubated at 60°C for 20 minutes to ensure the sample was completely dissolved. Samples were syringe filtered to remove any remaining undissolved protein before injecting 1 mL onto the Superdex 200 column.

SAXS data for CAHS gels were obtained by manually centering capillaries containing premade gels in the X-ray beam. Data was recorded as a series of thirty 0.2 second exposures, but only the first exposure was analyzed to minimize artifacts from X-ray damage. The final analyzed data was corrected for scattering from the empty capillary and a buffer containing capillary. CAHS gel containing samples were made by dissolving 100 mg/mL lyophilized protein in a buffer containing 20 mM Tris pH 7 and 50 mM NaCl. The sample was incubated for 20 minutes at 60°C to ensure the protein was completely dissolved. Double open-ended quartz capillaries with an internal diameter of 1.5 mm (Charles Supper) were used to make the samples. Dissolved protein was directly drawn into the capillary via a syringe. Concentration gradients were generated by layering the protein with buffer. Both ends of the capillary were then sealed with epoxy. Samples were allowed to cool for 5 hours prior to measurement. Data were collected along the concentration gradient by collecting data in 2 mm increments vertically along the capillary.

All data analysis was done using custom MATLAB (Mathworks) scripts. First, an effective concentration was calculated by assuming the maximum concentration was 100 mg/mL and scaling the remaining samples by the integrated intensity of the form factor. It should be noted that the actual concentration could be significantly less than 100 mg/mL in the maximum concentration sample. Data was fit to an equation containing three elements to describe the features apparent in the scattering data. The high-angle form factor was modeled using a Lorentzian-like function to extract the correlation length and an effective fractal dimension.

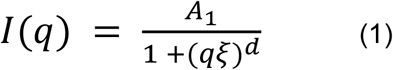

The correlation length is given by *ξ* and is related to the mesh size inside the fiber bundles seen in SEM images. The fractal dimension, *d*, is related to the density of the mesh. No clear correlation length was observed in the smallest angle data, and thus a power law was used to account for this component. The exponent *d*, is related to the nature of the interface inside and outside of the bundles.

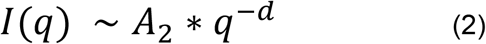

Finally, a Lorentzian peak was used to fit the diffraction peak that is apparent at higher concentrations. The width of the peak, B, appeared constant and was thus fixed so that the amplitude could be accurately extracted.

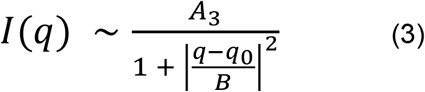

In all fit components, A_x_ is a scale factor.

### CD Spectroscopy

Lyophilized protein constructs were weighed and dissolved in a 20 mM Tris-HCl (Fisher Bioreagents, Hampton, NH) buffer at pH 7.0. CD spectra were measured using a JASCO J-1500 CD spectrometer with 0.1 cm quartz cell (Starna Cells, Inc, Atascadero, CA) using a 0.1 nm step size, a bandwidth of 1 nm, and a scan speed of 200 nm/min. Each spectrum was measured 7 times and averaged to increase signal to noise ratio. The buffer control spectrum was subtracted from each protein spectrum. Each protein construct was measured at several concentrations to ensure there is no concentration-dependent change in CD spectra (Figure 1 – figure supplement 1b, lower panels).

The resulting spectra were analyzed using the lsq_linear function from the SciPy library. To do this, base spectra for α-helix, β-sheet, and random coil spectra (taken from [Perczel, Park, and Fasman 1992] and available in Figure 1 – figure supplement 1b, upper right panel) were linearly fit to match the experimental data set. The three fit coefficients were normalized to give the relative contribution of each base spectrum to the experimental spectrum.

### Thioflavin T Assay

Proteins were dissolved in phosphate buffered saline, pH 7.2 (Sigma-Aldrich) was prepared in Dimethyl Sulfoxide (Sigma -Aldrich), and diluted to 20 μM in PBS for use in the assay. Thioflavin T was prepared fresh for each assay at a concentration of 400 μM in DMSO, then diluted to 20 μM in PBS. Twenty five microliters of protein and ThT in PBS were combined in a 96-well plate (Costar, Fisher Scientific, Hampton, NH) and incubated for 15 minutes at room temperature in the dark. Fluorescence was measured using a plate reader (Spark 10M, Tecan, Männedorf, Switzerland) with an excitation at 440 nm, emission was collected at 486 nm. CLC was suspended in 20 mM Tris buffer pH 7.5, the Thioflavin T was diluted in this buffer as well when using CLC. All proteins and controls were assayed in triplicate.

### Lactate Dehydrogenase Protection Assay

LDH desiccation protection assays were performed in triplicate as described previously.(Boothby et al., 2017) Briefly, protectants were resuspended in a concentration range from 20 g/L to 0.1 g/L in 100 μL resuspension buffer (25 mM Tris, pH 7.0). Rabbit Muscle L-LDH (Sigma-Aldrich) was added to this at 0.1 g/L. Half each sample was stored at 4°C, and the other half was desiccated for 17 hours without heating (OFP400, Thermo Fisher Scientific, Waltham, MA). Following desiccation all samples were brought to a volume of 250 μL with water. The enzyme/protectant mixture was added 1:10 to assay buffer (100 mM Sodium Phosphate, 2 mM Sodium Pyruvate [Sigma-Aldrich], 1 mM NADH [Sigma-Aldrich], pH 6). Enzyme kinetics were measured by NAD+ absorbance at 340 nm, on the NanodropOne (Thermo Fisher Scientific). The protective capacity was calculated as a ratio of NAD+ absorbance in desiccated samples normalized to non-desiccated controls.

### Citrate Synthase Protection Assay

The Citrate Synthase Kit (Sigma-Aldrich) was adapted for use in this assay (Chakrabortee et al., 2012; Goyal et al., 2005). All samples were prepared in triplicate, except desiccated negative control samples, which were prepared in quadruplicate, so that the extra sample could be used for assessment of desiccation efficiency. Concentration of gelatin (Sigma-Aldrich) was determined based on an average mass of 150 kDa. Lyophilized variants were resuspended in either purified water (samples to be desiccated) or 1X assay buffer (control samples) to a concentration of 20 g/L and diluted as necessary for lower concentrations. Citrate synthase (Sigma-Aldrich) was added at a ratio of 1:10 to resuspended protectants. Non-desiccated control samples were measured as described in the assay kit immediately following resuspension. Desiccated samples were subjected to 5-6 rounds of desiccation and rehydration (1-hour speedvac desiccation [Thermo Fisher Scientific] followed by resuspension in water). Following the 5^th^ round of desiccation, a negative control sample was resuspended and assayed to determine if activity remained. If the negative control sample retained more than 10% activity, a 6^th^ round of desiccation/rehydration was performed. After the final round of desiccation, samples were resuspended in 10 μL of cold 1X assay buffer. Samples were diluted 1:100 in the assay reaction mixture supplied, and all subsequent steps followed the kit instructions. The colorimetric reaction was measured for 90 seconds at 412 nm using the Spark 10M (Tecan).

### TD-NMR sample preparation

Quantitated and lyophilized protein samples were transferred as powder into 10 mm TD-NMR tubes (Wilmad Lab Glass) and resuspended in 500 μL of water to a final concentration of 1 g/L, 5 g/L, 10 g/L, 15g/L, and 20 g/L. Samples were left at room temp for 5 minutes to solubilize. If solubilization was not occurring (as determined visually), samples were moved to 55°C for intervals of 5 minutes until solubilized. If solubilization was not progressing at 55°C after 10 minutes of heating (as determined visually), then samples were transferred 95°C for 5 minutes intervals until fully solubilized. Samples were allowed to return to room temperature before being stored at 4°C. Measurements were taken within 72 hours following solubilization.

### Measurement of T_2_ relaxation

Low field Time-Domain NMR (TD-NMR) measures frequencies in the 10^9^ -10^12^ range, which is the frequency of ^1^H nuclei motion. Measurements in this frequency range allow for observation of water dynamics. Spin-spin relaxation (T_2_) is the measurement of ^1^H spins in a plane transverse to the magnetic field. T_2_ relaxation measured at low field strength is particularly useful for applications measuring slower water dynamics and water caging in porous structures (Valori et al., 2013; Zipp et al., 1976; Ectors et al., 2016; Emsley and Feeney, 1999) (Figure 5 – figure supplement 1a). T_2_ relaxation is dependent on the state of matter of the sample being measured, because state of matter impacts the ability of molecules to interact with one another. Water molecules in the bulk liquid state are free to interact with each other, and this interaction serves to hold the spins aligned for a longer period of time than water that is coordinated or caged, where the motion of water molecules is restricted (Figure 5 – figure supplement 1a). To better understand the impact of state of matter on spin decay, consider the analogy of hydrogen nuclei as magnets; two magnets that are free to interact will exert force on each other to maintain their magnetic alignments, whereas two magnets that are not free to interact influence their respective alignments less. Nuclei that are in the solid state or are heavily caged are essentially independent, because of their conformational rigidity and distance between them, and therefore their T_2_ relaxation is rapid. Nuclei in the bulk liquid state, on the other hand, can interact freely with other molecules in solution, and therefore their spin decay is slower.

These principles can also be applied to the ability of a protein to interact with water. For a theoretical protein to which water is blind, no interactions would occur and thus water molecules in this protein solution would behave as if it were a pure liquid, with a similar T_2_ relaxation profile. A highly hydrophobic protein, on the other hand, would exclude water molecules from itself, increasing the local density of water molecules and thus increasing their interactions, which would further slow T_2_ relaxation relative to bulk water. Hydrophilic proteins coordinate water molecules through electrostatic interactions, causing an ordering of water molecules into hydration shell(s) surrounding the protein. Being organized into hydration shells decreases the freedom of motion and interactions between different molecules of water, which would speed up T_2_ relaxation. Thus, T_2_ relaxation can inform us about the ability of a protein to coordinate or exclude water.

TD-NMR was performed using a Bruker mq20 minispec low-field nuclear magnetic resonance spectrophotometer, with a resonance frequency of 19.65 MHz. Samples were kept at 25°C during measurements through the use of a chiller (F12-MA, Julabo USA Inc., Allentown, PA) circulating a constant-temperature coolant. T_2_ free induction decays were measured using a Carr-Purcell-Meiboom-Grill (CPMG) pulse sequence with 8000 echoes, and an echo time of 1000 μs. Pulse separation of 1.5 ms, recycle delay of 3 ms, and 32 scans were used for all samples. Gain was determined for each sample individually and ranged from 53-56. Conversion of the free induction decay to T_2_ relaxation distribution was processed using the CONTIN ILT software provided by Bruker. Each variant was measured for the full concentration range, along with a buffer control (water or 20 mM Tris pH 7) in a single day.

### Measurement of Diffusion Coefficients

The diffusion coefficient was determined by the pulsed field gradient spin echo (PFGSE) method using a gradient pulse of 0.5 ms, gradient pulse separation 7.51416 ms, 90°–180° pulse separation 7.51081 ms and 90° first gradient pulse of 1 ms. Each variant was measured for the full concentration range, along with a buffer control (water or 20 mM Tris pH 7) on the same day that T_2_ relaxation data was collected.

In this work, diffusion coefficients of water molecules in protein solutions are represented relative to the diffusion coefficient of the water molecules in bulk solution, to emphasize the differences between proteins. This is calculated as: Buffer Diffusion minus Protein Diffusion, and is referred to as Δ Diffusion Coefficient (ΔDC).

### Analysis of Secondary Structures within CAHS D Glasses with Fourier Transform Infrared (FTIR) Spectroscopy

The kinetics of amide (peptide NH) proton exchange can in principle provide information on the flexibility of proteins. The slowing or inhibition of hydrogen/deuterium exchange can be attributed to either the involvement of the amide hydrogen in hydrogen bonding through secondary structure interactions, or to limited accessibility of the amide group to the deuterated solvent. It has been shown that a hydrogen bonded amide proton is protected from exchange even when it is exposed to solvent, slowing down exchange rates by more than six orders of magnitude (Barth, 2007). On the other hand, solvent exclusion can retard the exchange by a similar extent. Specific amide groups can be accessible to the solvent only in conformational substates that are rarely populated in the time average structure. Hydrogen exchange can therefore be extremely sensitive to the protein conformational dynamics. We previously assessed the impact of solvent accessibility (Figure 1e, Figure 3 - supplement 1c-e), and here we use an isopiestic approach to determine how hydrogen-bonding impacts the extent and the kinetics of hydrogen/deuterium exchange in CAHS glassy matrices at different hydration levels.

### Analysis of the amide I’ band in CAHS gels and glasses at increasing hydration

The amide I band centered around 1650 cm^-1^ is sensitive to the structure of the protein backbone; as a result a particular secondary structure absorbs predominantly in a specific range of the amide I region (Barth, 2007). However the various secondary-structure components overlap leading to a rather broad and scarcely structured band and band-narrowing procedures are necessary to resolve the component bands (Barth and Zscherp, 2002). The linewidth of the second and fourth derivative of a band is smaller than that of the original band, therefore the minima and maxima of these derivatives can be used to evaluate the number and positions of overlapping spectral components (Butler, 1970). On this basis, information on the secondary structure of the protein can be gleaned by fitting the amide I band to the sum of its component derivative bands. The component bands are assigned to specific secondary structures, and the fractional area of each component band is taken as the relative content of the corresponding secondary structures. We have used this approach with the aim of assessing CAHS secondary structure and its possible changes upon vitrification and dehydration.

### FTIR Sample Preparation

All samples were prepared by dissolving the CAHS protein in D_2_O to avoid the overlapping of the spectral contribution due to the H_2_O bending mode (around 1640 cm^-1^) (Maréchal, 2011) with amide I band of the CAHS protein centered at approximately 1650 cm^-1^. Deuteration causes a blue shift of the water bending mode by more than 400 cm^-1^, while inducing only a small shift of the amide I band (amide I’ band). Gel samples were obtained by dissolving the lyophilized CAHS protein at a concentration of 16 g/L in D_2_O heated at about 50 °C and gently stirred for 5 minutes. A volume of 10 μL of the CAHS solution was deposited between two CaF_2_ windows, separated by a 50 μm teflon spacer, and mounted into a Jasco MagCELL sample holder. When cooled at room temperature (25°C) CAHS solution forms a gel.

Glassy samples were obtained by depositing a volume of 38 μL of the heated CAHS solution (16 g/L) in D_2_O on a 50 mm diameter CaF_2_ window. Before gelling at room temperature, the drop spreads out to form a layer of approximately 1 cm diameter. The sample was dried under N_2_ flow for 5 minutes and subsequently the optical window on which the glass had formed was inserted into a specifically designed sample holder equipped with a second CaF_2_ window to form a gas-tight cylindrical cavity (Malferrari et al., 2011). Different hydration levels of the CAHS glass were obtained by an isopiestic method, *i*.*e*., by equilibrating the sample with saturated salt solutions providing defined values of relative humidity, contained at the bottom of the gas-tight sample holder cavity. The following saturated solutions in D_2_O were employed to obtain the desired relative humidity (Greenspan, 1977) at 297 K; KNO_3_ (RH=95%), NaCl (RH=75%), K_2_CO_3_ (RH=43%), and LiCl (RH=11%).

FTIR absorption measurements were performed at room temperature with a Jasco Fourier transform 6100 spectrometer equipped with a DLATGS detector. The spectra were acquired with a resolution of 2 cm^−1^ in the whole mid-IR range (7,000 - 1,000 cm^−1^) using a standard high-intensity ceramic source and a Ge/KBr beam splitter. All spectra were obtained by averaging 10^3^ interferograms.

### Amide I’ Band Analysis

Spectral analysis of the amide I bands recorded in gel samples was performed after subtracting a normalized D_2_O spectrum. Because of the low residual D_2_O in glassy samples, this procedure did not result in significant corrections in the amide band region.

When extracting the amide I’ band from the spectrum, the background was approximated by a straight baseline drawn between the minima on either side of the band. Second and fourth derivative spectra in the amide I’ region were calculated using the *i-signal* program (version 2.72) included in the SPECTRUM suite (http://terpconnect.umd.edu/~toh/spectrum/iSignal.html) written in MATLAB language. A Savitsky-Golay algorithm was employed to smooth the signals and calculate the derivatives.

Smoothing width was manually determined for each sample by optimizing the signal to noise ratio, by determining the impact of smoothing width on the calculated derivative spectra with the requirement of highest S:N without loss of spectral information. For all the amide I’ bands analyzed, in gel and glassy samples, both the second and fourth derivative spectra (not shown) were consistently dominated by three minima and maxima, respectively, suggesting the presence of three spectral components. This is supported by the finding that the positions of each peak detected in the second and fourth derivative spectra for each amide I’ band analyzed were very close (in most cases coincident within our spectral resolution).

The peak wavenumbers inferred from this analysis were found in the intervals 1619-1628 cm^-1^, 1644-1648 cm^-1^, and 1676-1688 cm^-1^. On this basis, amide I’ bands were decomposed into three Gaussian components by using a locally developed least-squares minimization routine (Malferrari et al., 2015) that applies a modified grid search algorithm (Bevington and Bevington, 1969). Confidence intervals for the best-fit parameters were evaluated numerically, as previously detailed (Francia et al., 2009). In the fitting procedure, the peak wavenumbers of the three Gaussian components were allowed to be varied over the intervals reported above, while the areas and the widths were treated as free, unconstrained parameters.

### Measurement of conformational dynamics of photosynthetic reaction centers embedded in CAHS glasses

The photosynthetic reaction center (RC) from the purple bacterium *Rhodobacter* (*Rb*.) *sphaeroides* represents an ideal model system to probe matrix effects on conformational protein dynamics. This membrane-spanning pigment-protein complex catalyzes the primary photochemical events of bacterial photosynthesis. Following absorption of a photon, the primary electron donor (P) of the reaction center, which is a bacteriochlorophyll dimer situated near the periplasmic side of the protein, delivers an electron to the primary quinone acceptor, Q_A_, located 25 Å away from P and closer to the cytoplasmic side of the RC. This electron transfer process, occurring in about 200 ps, generates the primary charge separated state, P^+^Q_A_^-^. In the absence of the secondary quinone acceptor bound at the Q_B_ site (or in the presence of inhibitors which block electron transfer from Q_A_^-^ to Q_B_), the electron on Q_A_^-^ recombines with the hole on P^+^ by direct electron tunneling (Feher et al., 1989).

The kinetics of P^+^Q_A_^-^ recombination after a short (nanosecond) flash of light provides an endogenous probe of the RC conformational dynamics. In fact, in room temperature solutions, following light-induced P^+^Q_A_^-^ charge separation, the RC protein undergoes a dielectric relaxation from a dark-adapted to a light-adapted conformation, which stabilizes thermodynamically the P^+^Q_A_^-^ state (lifetime for charge recombination of about 100 ms). When this conformational relaxation is inhibited (by freezing the RC to cryogenic temperature in the dark (McMahon et al., 1998), or by dehydrating within trehalose glasses (Palazzo et al., 2002)) the recombination kinetics are accelerated (lifetime of about 20 ms) and become strongly distributed, mirroring the immobilization of the protein over a large ensemble of conformational substates. We describe the charge recombination kinetics of confined RCs by using a continuous distribution *p(k)* of rate constants *k*: (Palazzo et al., 2002)

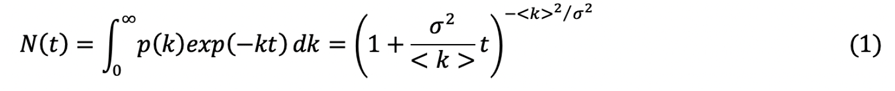

where *N(t)* is the survival probability of the P^+^Q_A_^-^ state at time *t* after the photo-activating light pulse, <k> is the average rate constant, and σ is the width of the rate distribution. The latter two parameters provide quantitative information on the conformational dynamics of the RC on the time scale of milliseconds. An increase in either or both parameters (<k>, σ) reflects a slowing of RC relaxation from the dark-to the light-adapted conformation (<k>), or of the fluctuation between conformational substates (σ) (Malferrari et al., 2015; Palazzo et al., 2002).

### Sample Preparation

*Seventy eight microliters* of RCs purified from *Rb. sphaeroides* R26 at 76 μM concentration in assay buffer (10 mM Tris, 0.025% LDAO, pH 8.0) was mixed with 64 μL of 16 g/L CAHS protein in water, and 8 μL of 200 mM o-phenanthroline in ethanol. O-phenanthroline is an inhibitor that blocks Q_A_^-^ to Q_B_ electron transfer, thus allowing the recombination kinetics of the P^+^Q_A_^-^ state to be monitored. The lyophilized CAHS protein was dissolved in water and heated to 50°C for 5 minutes. The protein was allowed to cool to room temperature, and during this cooling it was rapidly mixed with the RC suspension prior to gel formation. This mixture was immediately layered on a 50 mm diameter CaF_2_ optical window and dried under N_2_ flow for 5 minutes. The sample was then inserted into the gas-tight holder and equilibrated at a RH=11% in the presence of LiCl. The glassy matrix is characterized by a (CAHS/RC) molar ratio of approximately 6.6, corresponding to a mass ratio of about 1.7. This ratio was chosen for comparison with previous results of strongly inhibited RC conformational dynamics when embedded in glassy trehalose matrices (Malferrari et al., 2015).

### Time resolved optical absorption spectroscopy

The kinetics of P^+^Q_A_^-^ recombination was measured by time-resolved spectrophotometry, by recording the absorbance change at 422 nm following a 200 mJ pulse at 532 nm (7 ns width) provided by a frequency doubled Nd:YAG laser (Francia et al., 2008). In order to test the capability of the CAHS glassy matrix to inhibit RC dynamics, we measured P^+^Q_A_^-^ recombination kinetics in CAHS-RC glasses equilibrated at increasing RH using the isopiestic method described in “*FTIR Sample Preparation”*. At each relative humidity the residual water content of the CAHS-RC matrices was determined from the area of the (v_2_+v_3_) combination band of water at 5155 cm^-1^, using the absorption band of the RC at 802 as an internal standard (Malferrari et al., 2011; Malferrari et al., 2015). To this end, optical measurements were extended to the NIR region (15000-2200 cm^-1^) using a halogen lamp source, replacing the Ge/KBr with a Si/CaF2 beam splitter and the KRS-5 with a CaF_2_ exit interferometer window.

Since we do not know how the residual water is distributed between the different components which form the glassy matrix, evaluation of the residual water content allows for a physically meaningful comparison of the overall average hydration of the matrix between RCs embedded in either CAHS D or trehalose glasses (Figure 1 – figure supplement 1d). The water content, determined primarily as H_2_O:RC molar ratio, has been expressed as the ratio of mass water:mass dry matrix. To determine the mass of the dry matrix, we included the CAHS protein, the RC, and the detergent belt of the RC formed by 289 LDAO molecules^77^ plus 14 molecules of free LDAO per RC.

The hydration isotherms obtained using this approach in CAHS-RC and trehalose-RC glassy matrices indicate that the overall propensity for water adsorption is very similar in the disaccharide-RC and in the CAHS-RC protein matrix when the relative humidity is varied over a large range. Sorption data of the CAHS-RC matrix well fit the Hailwood and Horrobin equation (Hailwood and Horrobin, 1946).

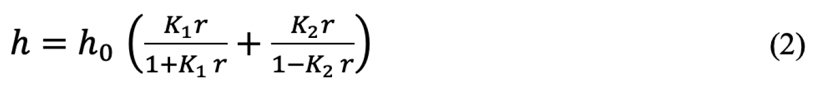

where h represents the equilibrium water content of the matrix, h_0_ and K_1_ are constants proportional to the number and activity of the hydration sites (respectively), and K_2_ is related to the water activity of the water condensing at the surface of the absorbing components. Best fitting to eq.2 yields h_0_=5.59 .10^−2^ g water/g dry matrix, K_1_=3.44 for the strong binding sites, and K_2_=1.04 for the weak condensing sites.

### Proteome-wide bioinformatics

The tardigrade proteome (tg.default.maker.proteins.fasta), taken from https://github.com/sujaikumar/tardigrade was used. The proteome file was pre-processed using protfasta (https://protfasta.readthedocs.io/), IDRs predicted with metapredict(Emenecker et al., 2021) and IDR kappa values calculated using localCIDER (Das and Pappu, 2013; Holehouse et al., 2017). Metapredict identified 35,511 discrete IDRs distributed across 39,532 proteins in the tardigrade proteome.

### Dimension estimates for LDH and CS

Radii of gyration for LDH and CS were calculated from the tetramer structure in PDB:1I10 and for the monomeric CS in PDB:3ENJ (Larson et al., 2009; Read et al., 2001).

### All-atom simulations

All-atom simulations were run using the ABSINTH implicit solvent model and CAMPARI Monte Carlo simulation engine (https://campari.sourceforge.net/). ABSINTH simulations were performed with the ion parameters derived by Mao et al., 2012. The combination of ABSINTH and CAMPARI has been used to generate experimentally-validated ensembles for a wide range of IDRs (Martin et al., 2020, 2016; Metskas and Rhoades, 2015).

Simulations were performed for the full-length CAHS D protein starting from a randomly generated non-overlapping starting state. Monte Carlo simulations update the conformational state of the protein using moves that perturb backbone dihedrals, and sidechain dihedrals, and rigid-body coordinates of all components (including explicit ions).

Ten independent simulations were run for 150,000,000 steps each in a simulation droplet with a radius of 284 Å at 310 K. The combined ensemble generated consists of 69,500 conformations, with ensemble average properties computed across the entire ensemble where reported.

Given the size of the protein, reliability with respect to residue-specific structural propensities is likely to be limited, such that general trends should be taken as a qualitative assessment, as opposed to a quantitative description. Simulations were analyzed using soursop (https://github.com/holehouse-lab/soursop) and MDTraj (McGibbon et al., 2015).

## Supplemental Files

File supplement 1: File containing the protein and nucleotide sequence for each protein variant used in this study.

Movie 1: Supplemental movie of CAHS D ensemble.

## Figure supplement legends

**Movie 1: CAHS D is predicted to exist in a dumbbell-like ensemble**. Monte Carlo simulation of predicted ensemble states of CAHS D. Source data for simulations can be found on GitHub (see Source Data section).

**Figure 1 – figure supplement 1:** Properties of hydrated and desiccated CAHS D gels. a) Concentration gradient gel SAXS structure probing revealed an emergent structure of approximately 9.5 nm. Areas indicated in the SAXS plot correspond with structures shown in SEM images, with the 9.5nm feature corresponding to approximate fiber sizes, while larger features correspond with the void spaces between gel fibers. Top right panel shows the derivation of sizes for each region of the SAXS plot. Lower right panel shows a projected relationship of the size of fibers with increasing protein concentration. b) CD spectra for C-terminus, CAHS D, Fl-Proline, Linker Region, and N-terminus. Top panels show the spectra for each variant listed at 20 or 25 μM (in green), overlayed with the resulting fit of the base spectra (dashed black line). Rightmost top panel shows the base spectra used to fit the experimental spectra (Perczel 1992). Lower panels show the concentration dependence of the spectra for each variant, at a range of concentrations. No structural changes are observed in the range tested. Premature rise at low wavelengths occuring at high concentrations in the CAHS D, FL-Proline, and Linker Region spectra are a result of reduced signal reaching the detector. Concentrations in μM are shown in the legend. c) Hydration dependent immobilization of biological material with the CAHS D glassy matrix. Charge recombination kinetics after a laser pulse (fired at t=0) are shown for RC incorporated into CAHS D glasses equilibrated at different relative humidity. Continuous curves represent best fit to eq.1. Reaction centers in solution (black line) and embedded into the CAHS D glass held 11% RH (purple line) have decay kinetics characterized by an average rate constant <k> = 8.8 s^-1^ and <k> = 35.5 s^-1^ respectively, and a rate constant distribution width of σ = 3.6 s^-1^ and σ = 23.8 s^-1^, respectively. With increasing relative humidity (23%-53% RH) the RC becomes more mobile. d) (Top) Hydration isotherms at 298 K determined in CAHS-RC (red circles) and trehalose-RC (black circles) amorphous matrices. In the glasses the CAHS/RC and trehalose/RC mass ratios were equal to 1.7. Continuous curve representing a best fit to eq. 2 of the water sorption data obtained in the CAHS-RC matrix (see supplemental text for details). Middle & lower panels show dependence of recombination kinetics on the residual water content in CAHS-RC matrices (red circles), in trehalose-RC glasses characterized by trehalose/RC molar ratios of 5000 and 500 (black squares and circles, respectively), and in a dehydrated RC film in the absence of any excipient (blue circles). Middle panel shows the average rate constant, <k>, bottom panel shows the rate constant distribution width, σ. Vertical bars represent confidence intervals within two standard deviations. The dashed lines give the value of <k> and σ obtained in solution. Dotted lines indicate the confidence intervals within two standard deviations. e) T_2_ relaxation of lysozyme, gelatin, and CAHS D. Concentration-dependent increase in relaxation kinetics is observed for CAHS D indicating strong water coordination, while gelatin shows a shift in T_2_ relaxation only after gelation occurs (20g/L). Lysozyme, a globular non-gelling protein, does not show strong water coordination. Source data for Figure 1 – figure supplement 1 is available in files: Figure 1 - figure supplement 1a - Source Data 1.zip, Figure 1 - figure supplement 1b - Source Data 1.zip, Figure 1 - figure supplement 1c - Source Data 1.xlsx, Figure 1 - figure supplement 1d - Source Data 1 (upper panel).xlsx, Figure 1 - figure supplement 1d - Source Data 2 (middle panel).xlsx, Figure 1 - figure supplement 1d - Source Data 3 (lower panel).xlsx and Figure 1 - figure supplement 1e - Source Data 1.xlsx.

**Figure 2 – figure supplement 1:** Transient structural properties of CAHS D. a) Model of interconversion of disordered linker helix, gaining and losing transient α helices. b) Calculated fractional α-helix (left) and β-sheet (right) content of the wildtype CAHS D as determined from simulations. c) Curve-fitting of radius of gyration determined from simulations (red points) overlaid with SAXS experimental data (black line). d) Calculated kappa value (amino acid charge distribution) of all proteins in the tardigrade proteome. Kappa provides a metric to quantify the degree of charge patterning, such that sequences with a lower kappa value have a more even distribution of charged residues. As shown, CAHS D has some of the most evenly distributed charges of all proteins in the proteome. Source data for Figure 2 – figure supplement 1 is available on GitHub.

**Figure 3 – figure supplement 1:** a) Bioinformatic predictions of the secondary structure of WT CAHS D, with arrows indicating where prolines were inserted into the sequence to disrupt secondary structures. b) SEM images of CAHS D and FL-Proline, showing that FL-Proline does not form the reticular meshwork gel structure found in CAHS D. c) Hydration-dependent FTIR spectra for CAHS D glasses stored at different relative humidities. Spectra for all hydration levels are overlaid. d) Deconvolution of FTIR spectra shown in (c) for the 1725-1575 cm^-1^ range. Top panel shows the deconvolution for 11% RH, and the bottom panel shows the same for 95% RH. Individual Gaussian components are shown in blue, orange and green; the red curve represents the corresponding best fit e) Evolution of the infrared spectrum between 1350 and 1750 cm^-1^ of an extensively dehydrated CAHS glass as a function of the deuteration time. The CAHS glassy matrix, previously equilibrated at RH=6% in the presence of NaOH.H_2_O, was exposed at t=0 to the atmosphere of a saturated NaCl solution in D_2_O, at RH=75%. Spectra are normalized to the amplitude of the amide I/I’ band. Source data for Figure 3 – figure supplement 1 is available in files: Figure 3 - figure supplement 1c - Source Data 1.xlsx, Figure 3 - figure supplement 1d - Source Data 1 (upper panel).xlsx, Figure 3 - figure supplement 1d - Source Data 2 (lower panel).xlsx, Figure 3 - figure supplement 1e - Source Data 1.xlsx.

**Figure 4 – figure supplement 1:** Sigmoidal curves representing the full range of protection for each variant using the LDH unfolding assay. b) Sigmoidal curves representing the full range of protection for each variant using the CS aggregation assay. Source data for Figure 4 – figure supplement 1 is available in files: Figure 4 - figure supplement 1a - Source Data 1.xlsx and Figure 4 - figure supplement 1b - Source Data 1.xlsx.

**Figure 5 – figure supplement 1:** CAHS variant water coordination. a) Example TD-NMR data and explanation. 1H nuclei transverse spin magnetization Free Induction Decays (FID) are subjected to an inverse Laplace transformation, which provides a distribution of spin-spin decay times referred to as T_2_ relaxation. The leftward shift of T_2_ peak midpoints indicates how water is ordered in the sample being measured. Bulk liquids (free water) have slower T_2_ relaxation than samples which display water coordination. This figure shows example T_2_ distributions, with representative illustrations of the water behavior in each sample. b) T_2_ relaxation distributions for all variants. c) Delta-diffusion coefficient *versus* concentration for all variants. Source data for Figure 5 – figure supplement 1 is available in files: Figure 5 - figure supplement 1b - Source Data 1.xlsx and Figure 5 - figure supplement 1c - Source Data 1.xlsx.

**Figure 5 – figure supplement 2:** a) Aggregation PD50 plotted against T2/MW for all variants at 1 g/L (top), 5 g/L (middle) and 20 g/L (bottom). No correlations are observed. b) Unfolding PD50 plotted against T2 peak midpoint for all variants at 1 g/L (top), 5 g/L (middle) and 20 g/L (bottom). Only very weak correlations are seen between non-normalized T2 and unfolding PD50.. c) Diffusion at 1mM plotted against PD50 for unfolding (left) and aggregation (right). d) Diffusion at 1mM normalized to MW, plotted against PD50 for unfolding (left) and aggregation (right). No strong correlations are found for diffusion and PD50. For parts (a)-(d) error bars represent standard deviation for PD50. Bars smaller than the size of the points are shown as white bars within points. Source data for Figure 5 – figure supplement 2 is available in files: Figure 5 - figure supplement 2a - Source data 1.xlsx, Figure 5 - figure supplement 2b - Source data 1.xlsx, Figure 5 - figure supplement 2c - Source data 1.xlsx and Figure 5 - figure supplement 2d - Source data 1.xlsx.

**Figure 6 – figure supplement 1:** Working models for CAHS D variant behavior and prevention of protein unfolding in the (re)hydrated, drying, and dry state. Schematics representation interpreted modes and degree of protection and gelation for each of our variants is presented.

**Figure 6 – figure supplement 2:** Working models for CAHS D variant behavior and prevention of protein aggregation in the (re)hydrated, drying, and dry state. Schematics representation interpreted modes and degree of protection and gelation for each of our variants is presented.

## Source Data

Figure 1d – Source Data 1.xlsx – Source data for CAHS, Gelatin, and Lysozyme delta diffusion coefficients.

Figure 1e – Source Data 1.xlsx – Source data for hydrogen deuterium exchange kinetics.

Figure 2d – Source Data 1.xlsx – Source data for circular dichroism spectroscopy secondary structure determinations.

Figure 2e – Source Data 1.zip – Source data for SAXS experiments.

Figure 3b - Source Data 1.xlsx – Source data for FTIR experiments on dry CAHS D protein

Figure 3c - Source Data 1.xlsx – Source data for thioflavin T labeling.

Figure 4a - Source Data 1.xlsx – Source data for PD50 data for LDH unfolding assay.

Figure 4b - Source Data 1.xlsx – Source data for PD50 data for CS aggregation assay.

Figure 4d - Source data 1.xlsx – Source data for delta diffusion coefficient data at 1mM for all variants.

Figure 5a - Source data 1.xlsx – Source data for TD-NMR for CASH D, 2x Linker, NLN, N- terminus, FL-Proline, and CLC variants.

Figure 5c - Source data 1.xlsx – Source data for unfolding assay PD50 vs T2/MW plot. Figure 5d - Source data 1.xlsx – Source data for aggregation assay PD50 vs T2/MW plot.

Figure 5e - Source data 1.xlsx – Source data for single variable plots (LDH PD50, T2/MW, CS PD50, and delta-DC).

Source data for Movie 1 and simulations can be found on GitHub at: https://github.com/holehouse-lab/supportingdata/tree/master/2021/hesgrove_CAHS_2021

Figure 1 - figure supplement 1a - Source Data 1.zip

Figure 1 – figure supplement 1b – Source Data 1.zip: Source data for CD spectroscopy studies.

Figure 1 - figure supplement 1c - Source Data 1.xlsx: Source data for reaction center kinetics.

Figure 1 - figure supplement 1d - Source Data 1 (upper panel).xlsx: Source data for water content versus relative humidity.

Figure 1 - figure supplement 1d - Source Data 2 (middle panel).xlsx: Source data for RC kinetics versus water content.

Figure 1 - figure supplement 1d - Source Data 3 (lower panel).xlsx: Source data for RC kinetics versus water content.

Figure 1 - figure supplement 1e - Source Data 1.xlsx: Source data for TD-NMR for CAHS D, lysozyme, and gelatin.

Figure 3 - figure supplement 1c - Source Data 1.xlsx: Source data for FTIR on CAHS D at various relative humidities.

Figure 3 - figure supplement 1d - Source Data 1 (upper panel).xlsx: Source data for CAHS D FTIR at 11% relative humidity.

Figure 3 - figure supplement 1d - Source Data 2 (lower panel).xlsx: Source data for CAHS D FTIR at 95% relative humidity.

Figure 3 - figure supplement 1e - Source Data 1.xlsx: Source data for time dependence on absorbance of CAHS D.

Figure 4 - figure supplement 1a - Source Data 1.xlsx: Source data for sigmoidal protection curves for LDH unfolding assay.

Figure 4 - figure supplement 1b - Source Data 1.xlsx: Source data for sigmoidal protection curves for CS aggregation assay.

Figure 5 - figure supplement 1b - Source Data 1.xlsx: Source data for TD-NMR for all variants.

Figure 5 - figure supplement 1c - Source Data 1.xlsx: Source data for delta-diffusion coefficients vs. concentration for all variants.

Figure 5 - figure supplement 2a - Source data 1.xlsx: Source data for aggregation vs T2 per kDA.

Figure 5 - figure supplement 2b - Source data 1.xlsx: Source data for unfolding vs T2 per kDA.

Figure 5 - figure supplement 2c - Source data 1.xlsx: Source data for unfolding and aggregation PD50 vs diffusion at 1mM.

Figure 5 - figure supplement 2d - Source data 1.xlsx: Source data for Unfolding and aggregation PD50 vs diffusion at 1mM/M

