## Supplemental Figures for "Tardigrade CAHS Proteins Act as Molecular Swiss Army Knives to Mediate Desiccation Tolerance Through Multiple Mechanisms"

Figure 1- Supplement 1.

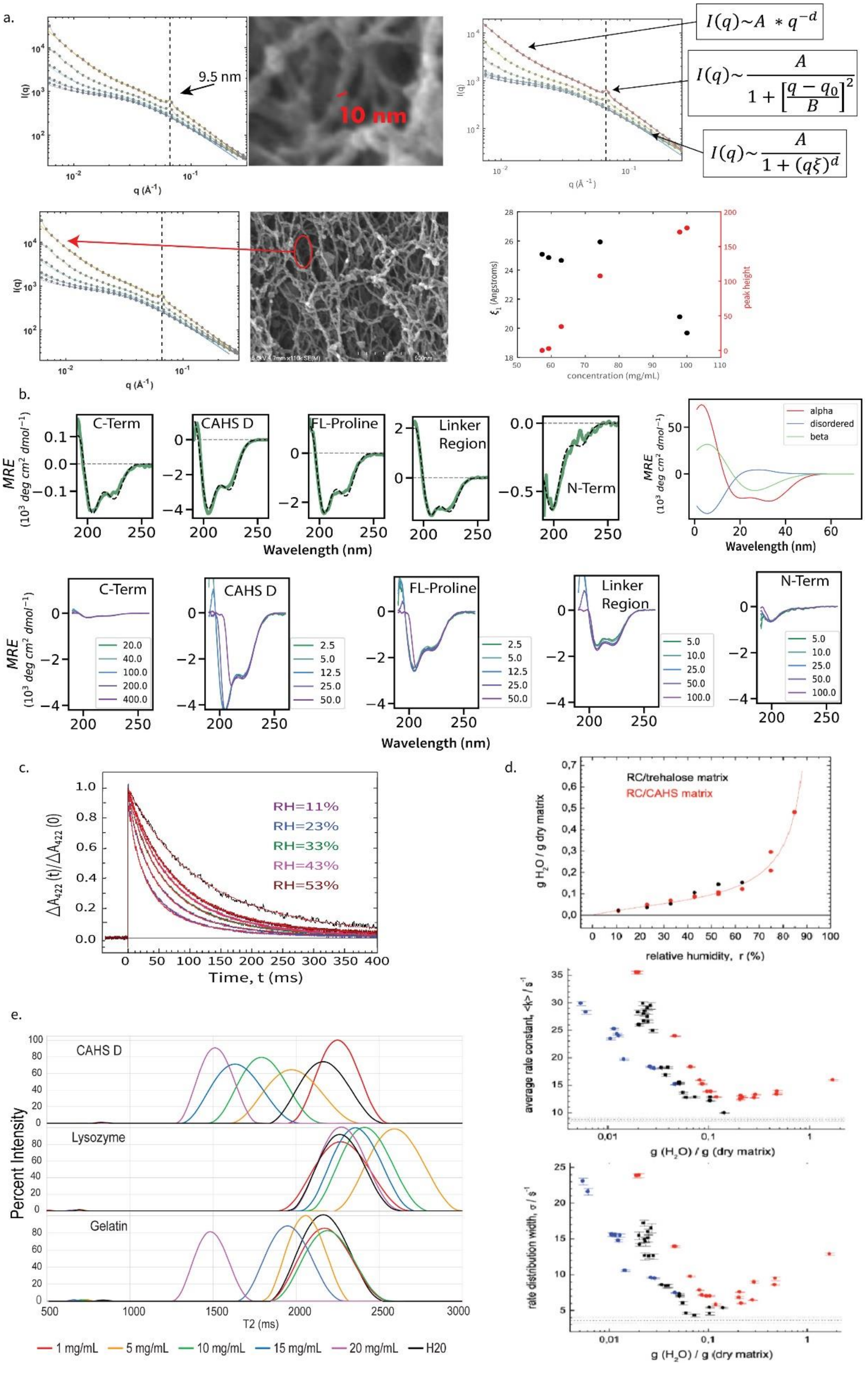

Figure 2- Supplement 1

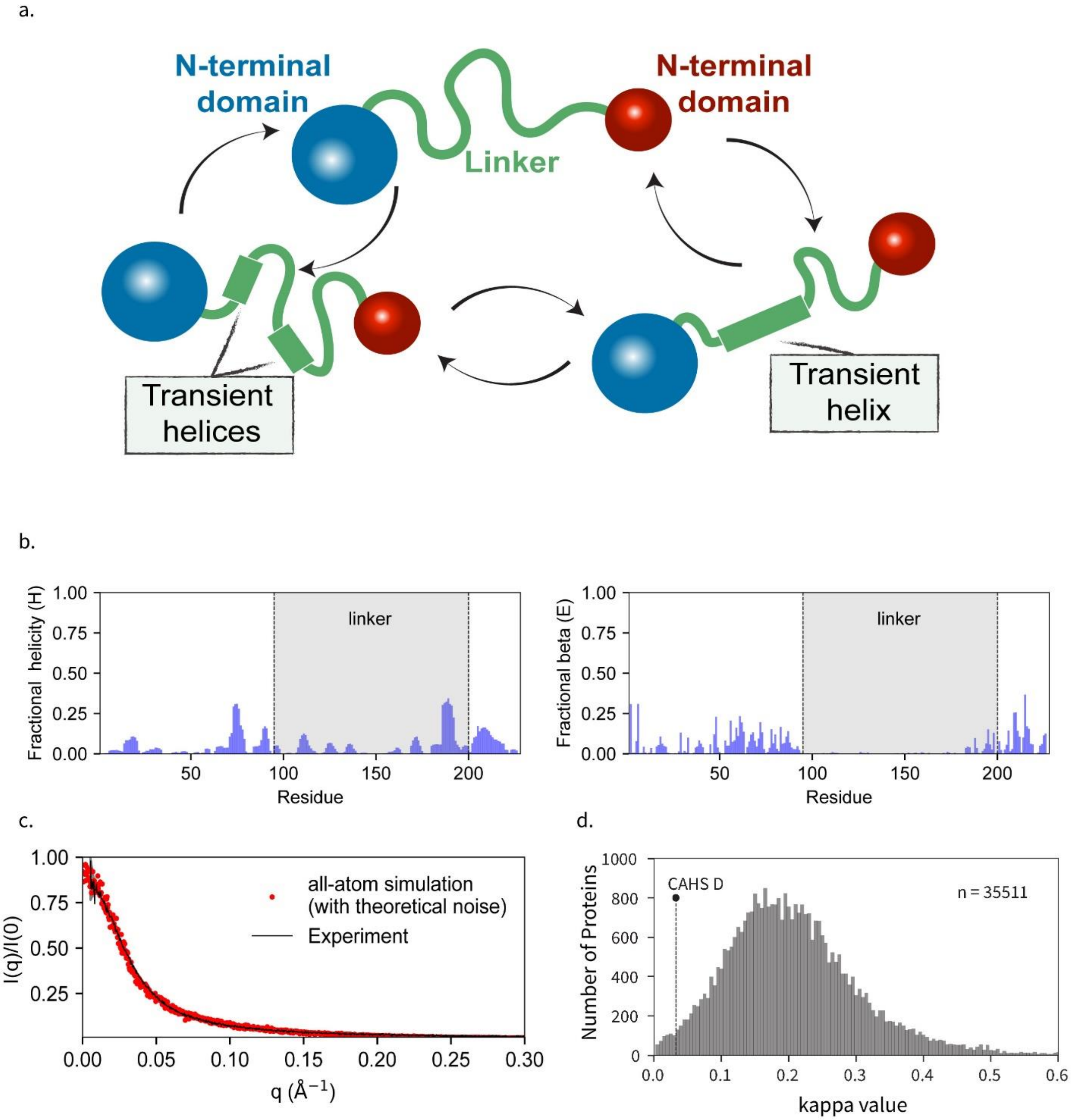

Figure 3- Supplement 1

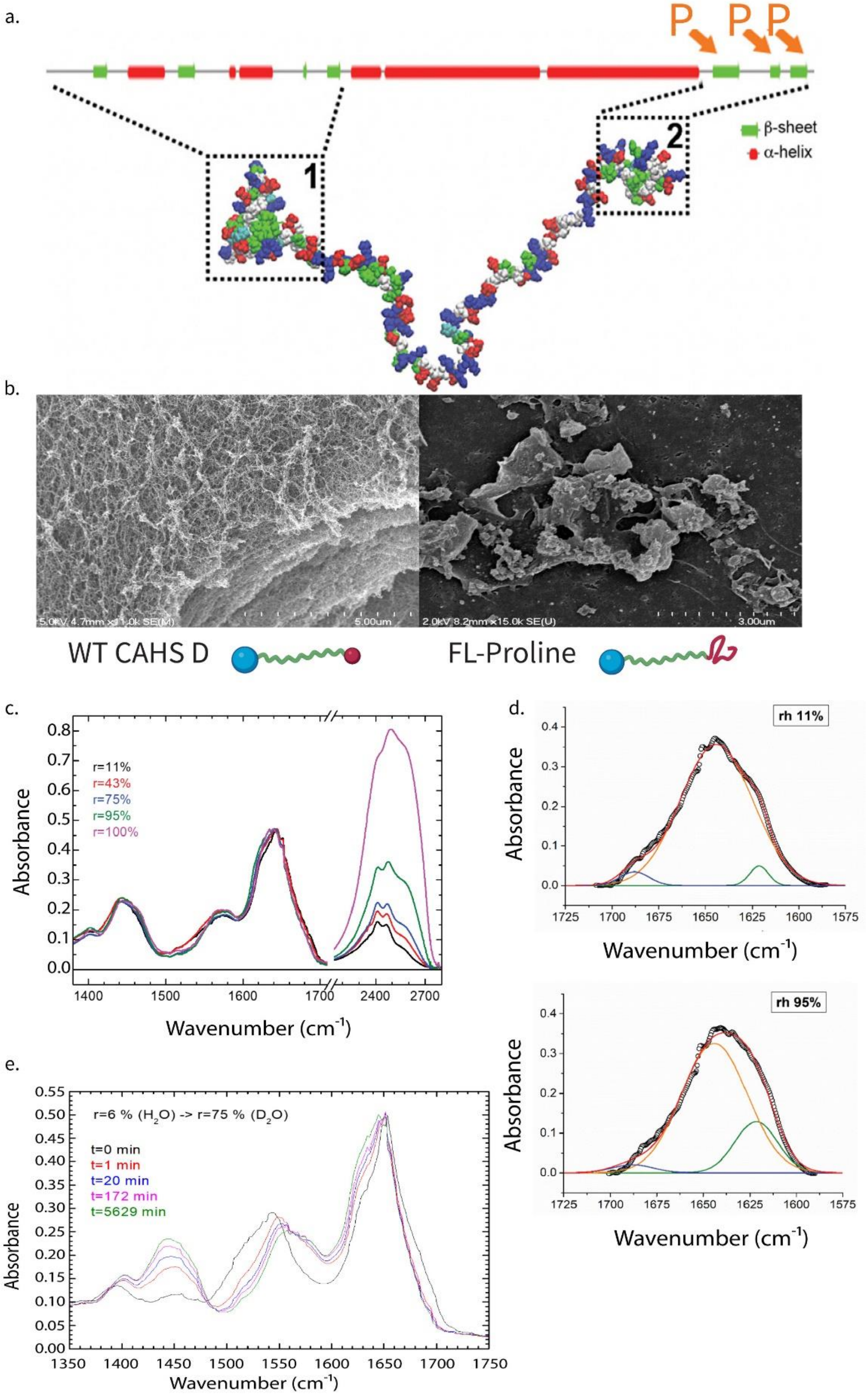

Figure 4- Supplement 1

a.

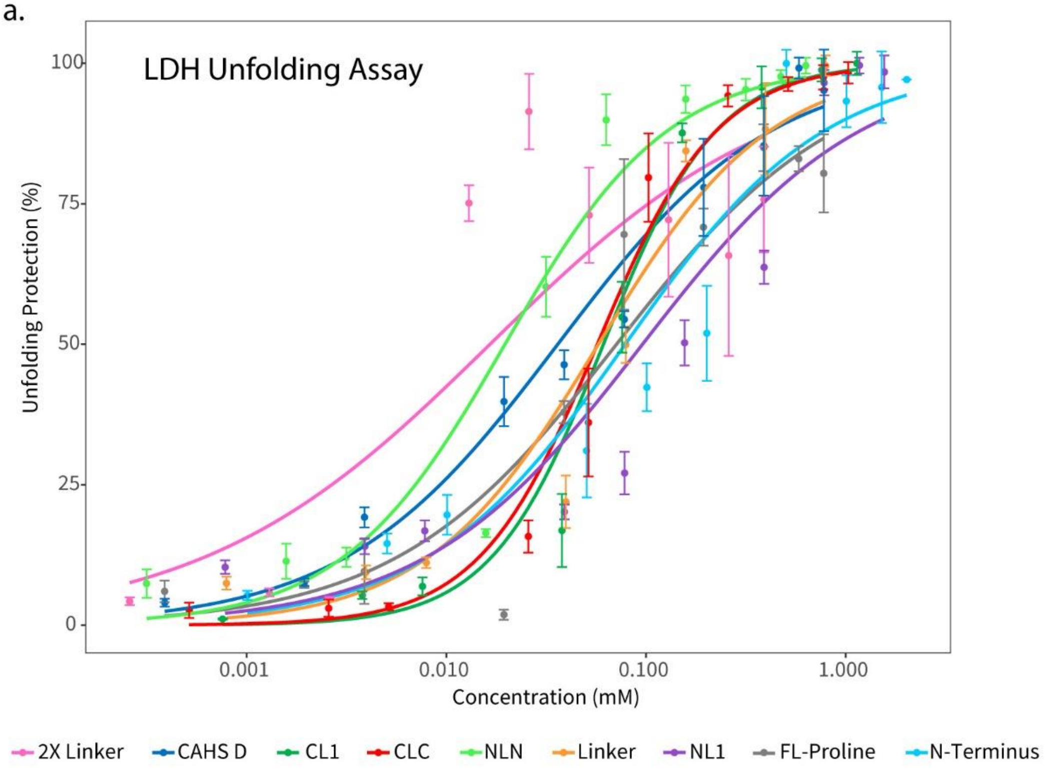


b.

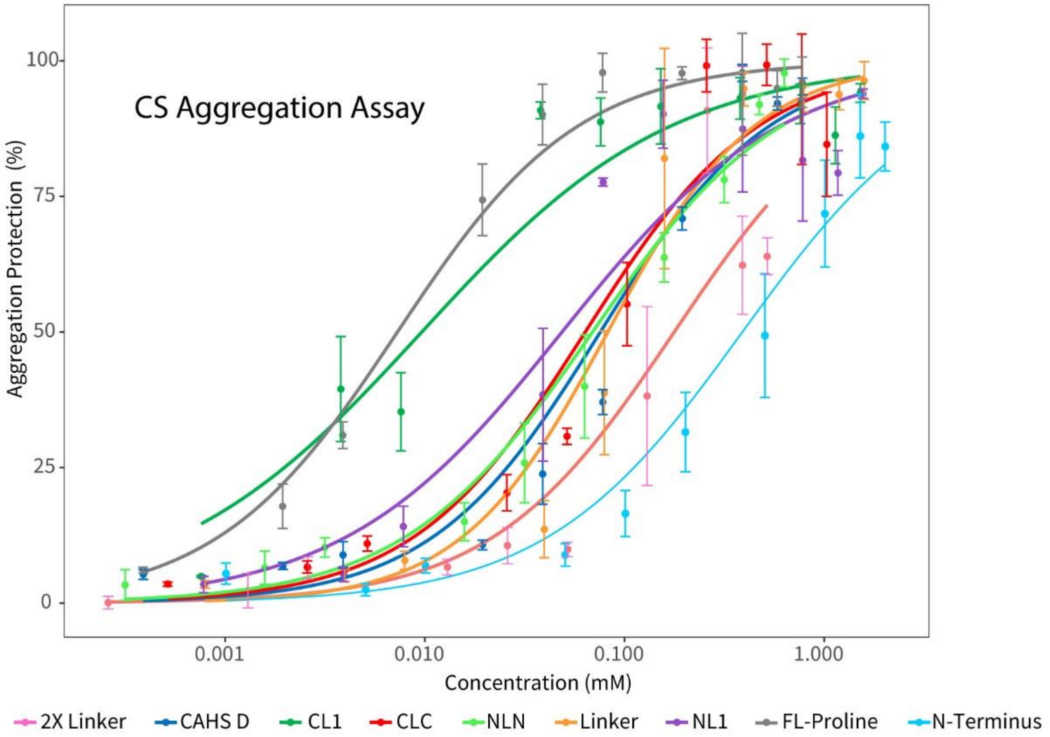

Figure 5- Supplement 1

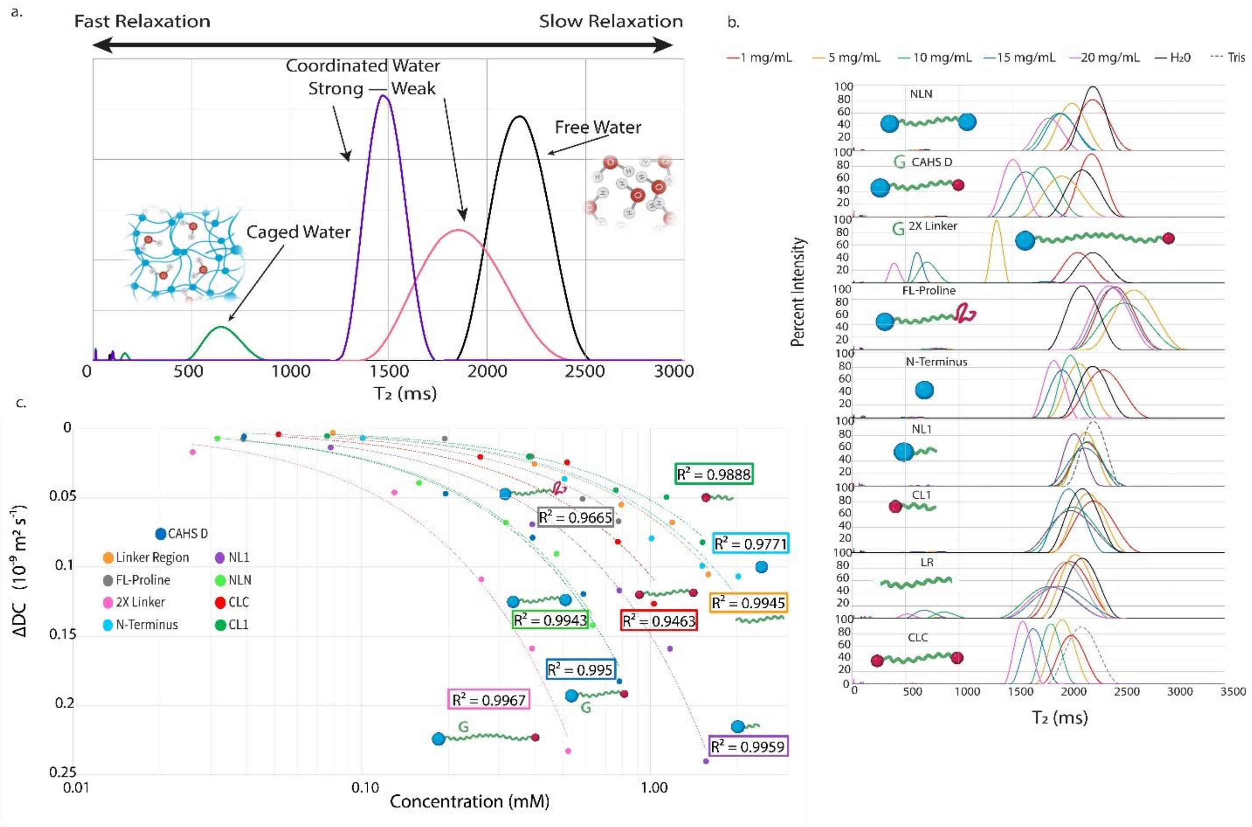

Figure 5- Supplement 2

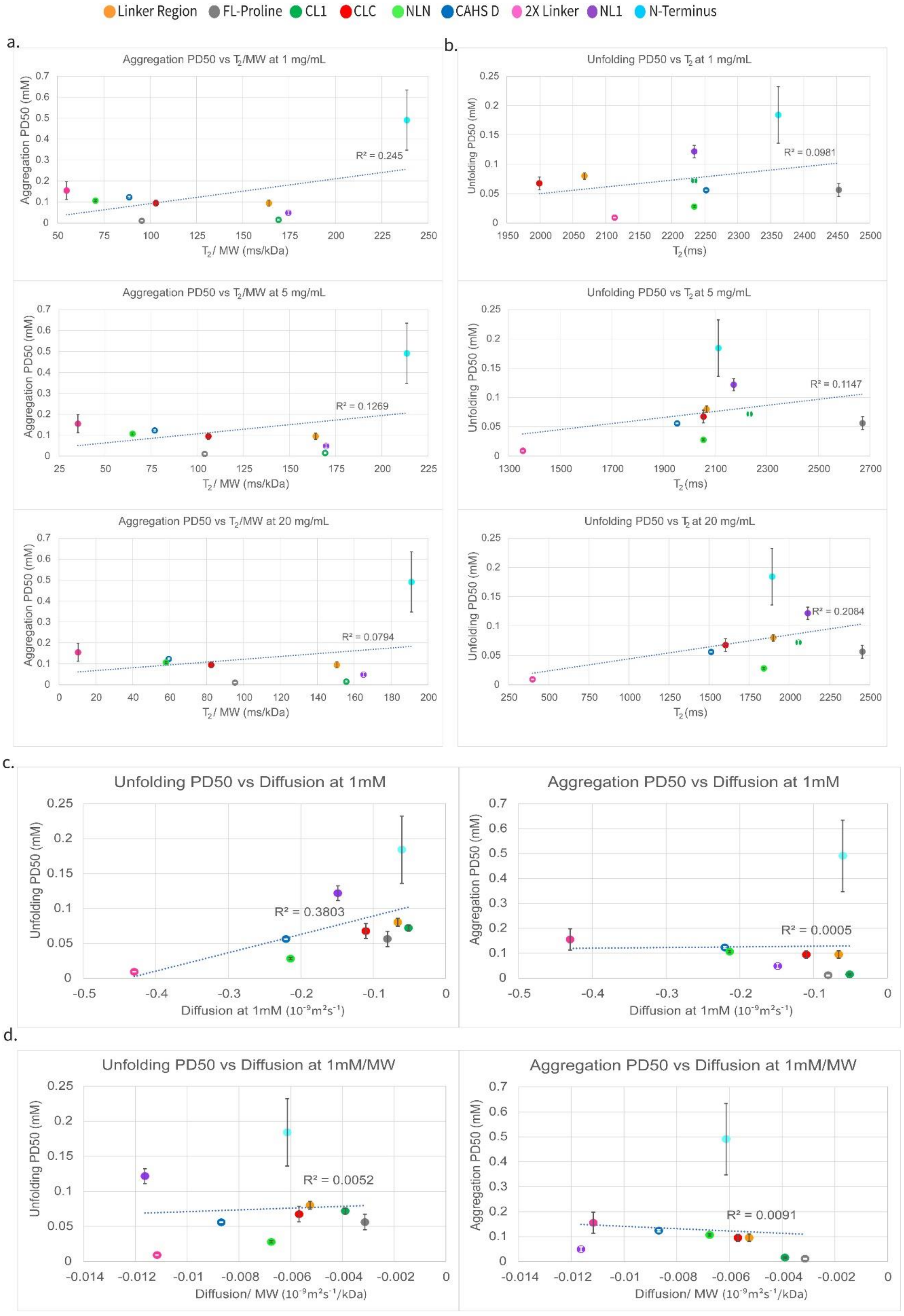

Figure 6- Supplement 1

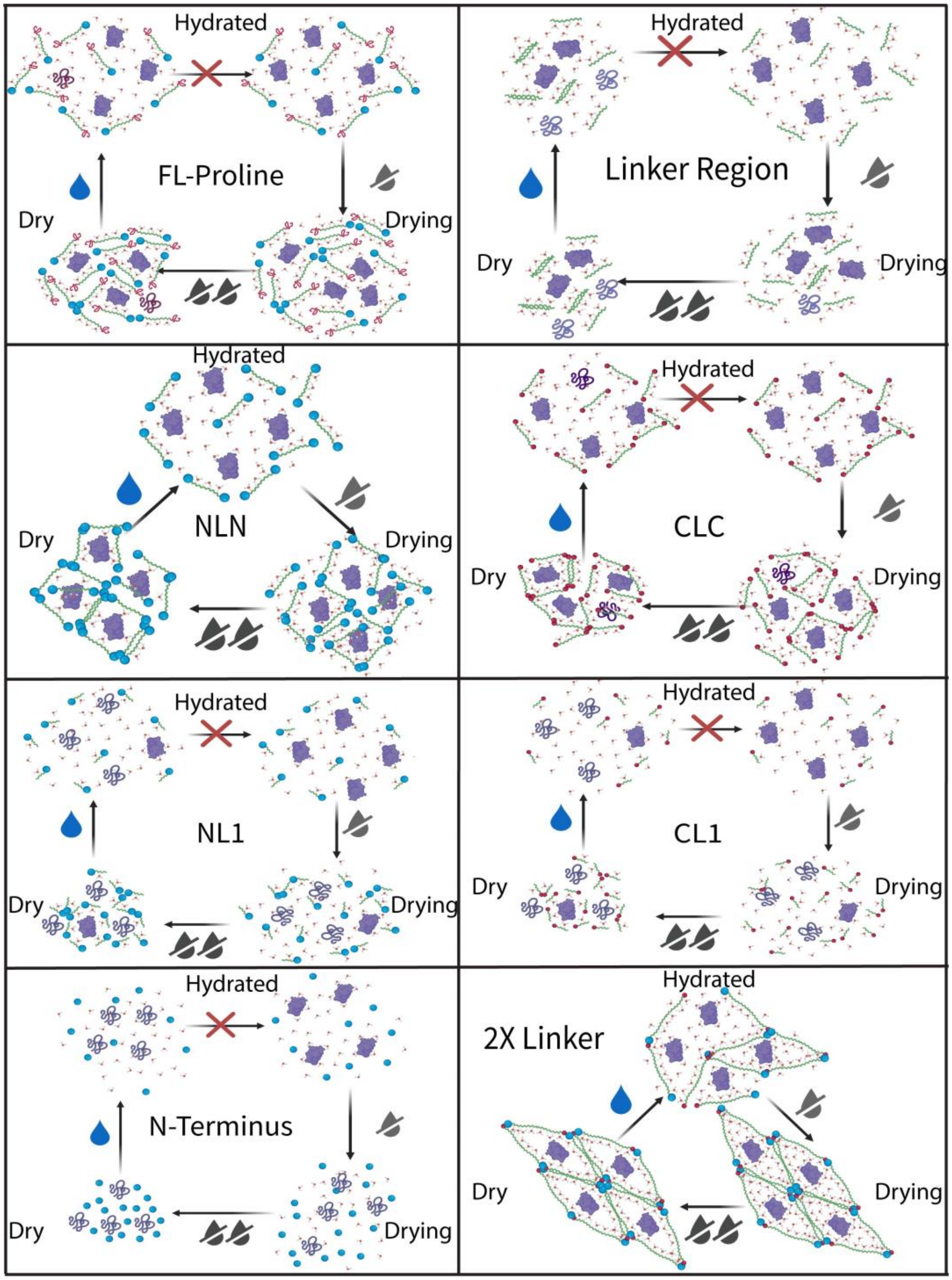

Figure 6- Supplement 2

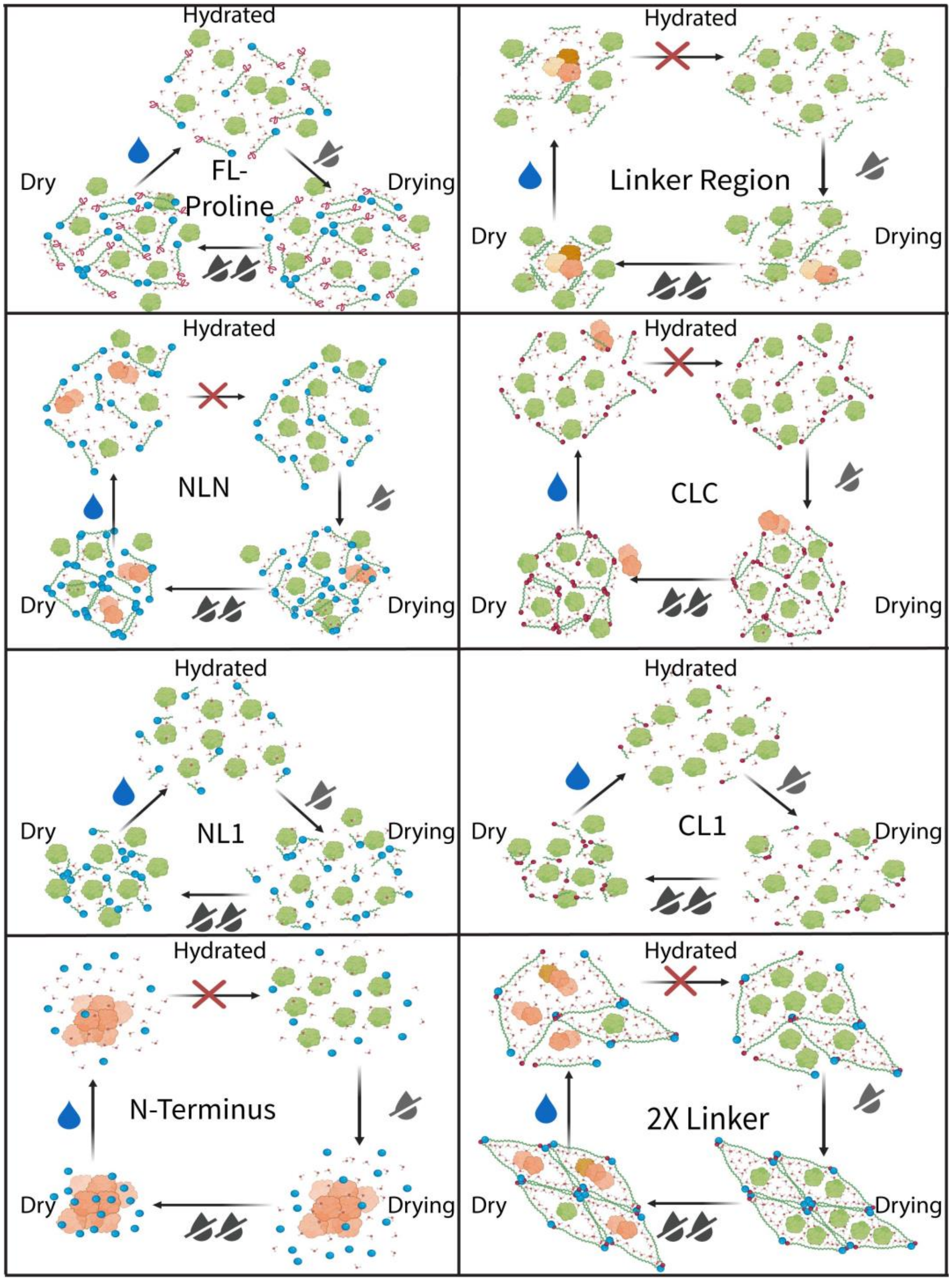
